# Lamins regulate nuclear mechanics and shape to control glioblastoma cell proliferation, migration and invasion

**DOI:** 10.1101/2024.09.30.615792

**Authors:** Xiuyu Wang, David Pereira, Isabelle Perfettini, Florent Péglion, Ryszard Wimmers, Ananya Roy, Karin Forsberg-Nilsson, Alexandre Baffet, Sandrine Etienne-Manneville, Jean-Baptiste Manneville

**Affiliations:** Laboratoire Matières et Systèmes Complexes, Université Paris Cité, CNRS UMR7057, 10 Rue Alice Domon et Léonie Duquet, F-75013, Paris, France; Cell Polarity, Migration and Cancer Unit, Institut Pasteur, CNRS, UMR3691, F-75015, Paris, France; Institut Curie, PSL Research University, CNRS UMR144, Paris, F-75246 France; Department of Immunology, Genetics and Pathology, Science for Life Laboratory, Uppsala University, Uppsala, Sweden; Science for Life Laboratory, Uppsala University, Uppsala, Sweden

## Abstract

Glioblastoma (GBM) is known as the most aggressive brain tumor and is characterized by a high heterogeneity and a median patient survival time around 15 months. Phenotyping based on cell mechanics is increasingly recognized as a potential prognostic marker for tumor aggressiveness. We have previously shown that cell and nuclear mechanical properties vary between different grades of gliomas and may be used to differentiate between GBM of different aggressiveness. Here, we find that the levels of lamin proteins can serve as an indicator of GBM aggressiveness. In patient-derived GBM cell lines, we found that cells from different GBM express different lamin levels. Nuclear size correlates positively with the ratio between lamin A and lamin B1 while nuclear stiffness increases with the levels of both lamin A and lamin B1. A simple mechanical model suggests that lamin A and lamin B1 act like springs in series. We also show that cells proliferate faster in GBM cell lines expressing higher lamin A levels. Downregulating lamin A expression in these cells reverse the aggressive phenotype. In contrast with breast cancer cells for which reduced lamin A levels favor cell migration in a confined environment, increased levels of lamin A may facilitate the invasion of more aggressive GBM cell lines in the soft environment of the brain. Furthermore, since nuclear deformation is a hallmark of malignancy in cancer cells, our results suggest that nuclear shape and mechanics may serve as prognosis biomarkers for GBM.

## Introduction

Glioblastoma multiforme (GBM), recognized as the most malignant primary brain tumor with an extremely poor prognosis[1], has long relied on the Tumor-Node-Metastasis (TNM) staging classification as the primary parameter to gauge its degree of malignancy[2]. Despite its widespread use, the TNM staging system, with GBM typically categorized as stage IV, fails to capture the considerable individual variability among patients, resulting in disparate prognoses even among seemingly similar tumors[3]. Grade IV GBM is rare, with an age-adjusted incidence ranging from 4.67 to 5.73 cases per 100,000 individuals worldwide, but devastating, as there is no curative treatment available[4, 5]. Even for diagnosis, only a few biomarkers are routinely used in clinical practice[6]. From diagnosis, prognosis, and treatment, GBM represent a longstanding challenge for clinicians[7]. Although their precise origin is still debated, GBM are thought to originate from glial cells, glial precursors or neural stem cells [8, 9]. In the case of isocitrate dehydrogenase (IDH)-wildtype GBMs, ventricular and outer radial glial cells in the subventricular zone of the brain have been suggested to give rise to GBM cells[9].

Following developments in the field of mechanotransduction, more and more research emphasizes that mechanical properties of both cancer cells and their environment are critical during tumour development[10] and metastasis[11, 12]. Mechanosensing and mechanosignalling are altered in cancer cells, which induces a remodeling of the surrounding extracellular matrix (ECM) and changes in cell mechanics, in particular at the level of the nucleus. As a consequence, abnormal cell nuclear morphology has often been used as a hallmark of cancer[13, 14]. In the case of glioma brain tumours, nuclear shape has been used for tumour grading by histology for more than forty years[15]. Moreover, *in vitro* studies using GBM cell lines have shown that substrate rigidity promotes GBM invasive characteristics such as cell spreading, motility and migration, as well as proliferation[16, 17]. More recent evidence shows that glioma stiffness is associated with increased mechanosensing and a highly invasive mesenchymal phenotype[18] and leads to the enhanced therapeutic resistance of GBM cells[19]

The mechanical properties of the nucleus are thought to be critical during tumour cell invasion[20, 12]. Being the largest organelle in the cell, the nucleus contributes significantly to the cell mechanical properties. Because cancer cells have to migrate through confined environments during tumour progression and metastasis, a softer nucleus could allow cancer cells to deform more and invade surrounding tissues more efficiently[21, 22]. The mechanical properties of the nucleus are imparted by the nuclear envelope which separates the nucleus from the cytoplasm and the chromatin which is packed inside the nucleus. Nuclear envelope mechanics is controlled by the nuclear lamina, a complex network of lamin intermediate filaments constituted of lamins A/C and lamins B[23]. The nuclear lamina is a dense fibrillar network underlying the inner nuclear membrane, composed primarily of A- and B-type lamins, which form a two-layered structure [24]. Lamins provide mechanical support to the nucleus and play a crucial role in regulating nuclear shape, stiffness, and chromatin organization [25, 26, 27]. It is now becoming clear that structural proteins of the nuclear lamina not only maintain a well-defined nuclear architecture but also participate in a wide range of cellular processes. For instance, mutations in lamin nuclear proteins cause the so-called laminopathies[28] and dysregulation of their expression is typically associated with the onset and spread of cancer[29]. The levels of lamins A/C and B have also been associated with cell and tissue stiffness. An elevated ratio between the expression levels of lamin A/C and the expression level of lamin B has been associated with cell and tissue stiffness[30], while a decrease in lamin A/C levels facilitates migration of breast cancer cells through narrow pores *in vitro*[31].

Here, we demonstrate that the nuclear lamin proteins play a critical role in the pathophysiology of GBM. In contrast with other techniques which infer rather than directly measure nuclear mechanics, we use indentation of the nucleus in living patient-derived GBM cells by internalized microspheres trapped in optical tweezers to quantify the mechanical properties of the nucleus. We correlate nuclear mechanics, lamin levels and indicators of GBM aggressiveness and suggest that, in contrast to what was previously reported in other cancer types, GBM invasion is facilitated by increased nuclear stiffness.

## Materials and Methods

### Cell culture

Glioblastoma (GBM) patient-derived cells lines (U3008, U3009, U3013, U3017, U3021, U3031, U3039, U3047, U3065, U3088, and U3123) were acquired from The Human Glioblastoma Cell Culture Resource (HGCC, Uppsala University, Sweden, www.hgcc.se)[32], which has the necessary ethical agreements to collect GBM samples from informed patients. GBM cells were cultured in DMEM/F-12 (Gibco) and Neurobasal Medium (Gibco) in a 1:1 ratio, supplemented with B27 (1X, Gibco), Penicillin (1%), and Epidermal growth factor (EGF, 10 ng/ml) and Fibroblast growth factor (FGF, 10 ng/ml). EGF and FGF were added just before use of the culture medium. Cells were cultured on Matrigel-coated plate (Gibco, Geltrex, LDEV-Free) at a concentration of 33 *µ*g/ml. Cell culture medium was changed every 3 days. Cells were passaged when 90% confluency was reached. The passage ratio was between 1/10 and 1/3 depending of the cell line proliferation rate.

Human fetal tissue samples were collected with previous patient consent and in strict observance of legal and institutional ethical regulations. The protocol was approved by the French biomedical agency (Agence de la Biomédecine, approval number: PFS17-003). The dissociation and culture procedure is described in [33]. Human radial glial (RG) cells were cultured in DMEM/F-12, supplemented with Glucose (Sigma, 2.9mg/mL), Penicillin(1%), and Amphotericin B (250 ng/mL). B27(-vitamin A) (1X, Gibco), and growth factors EGF (20 ng/ml) and FGF (20 ng/ml) were added before use. Culture dishes were coated with poly-D-lysine (PDL) at 2 *µ*g/cm^2^ in Phosphate Buffered Saline (PBS) for 1 hour then with fibronectin at 1 *µ*g/cm^2^ for 1 hour.

### siRNA transfection

Lamin A (LMNA gene) downregulation was performed using siRNA duplexes specific for human LMNA (si-LaminA) purchased from Eurofins Genetics. The si-LaminA Sequence (5’→3’) was AGA AGG AGC UGG AGA AGA C. Luciferase was used as control. The si-Luciferase Sequence (5’→3’) was UAA GGC UAU GAA GAG AGA C. GBM cells were transfected at confluency with si-LaminA and si-Luciferase using Lipofectamine 3000 (Thermo Fisher). Experiments were performed 72 hours after transfection.

### Western Blotting

Cells were lysed in RIPA buffer (Thermo Fisher) supplemented with Protease inhibitor (Sigma) on ice for 10 minutes, then collected and centrifuged at maximum speed 15200 rpm at 4 °C for 10 minutes. Protein concentration was determined using the Bradford assay (Bio-Rad Laboratories). Equal amounts of protein were mixed with reducing NuPAGE loading buffer (Invitrogen), boiled and electrophoresed on NuPAGE gels (Invitrogen), and then transferred to the membrane using iBlot 3 Transfer Stacks (Invitrogen). Blocking was performed for 1 hour with 5% nonfat dry milk in TBST and blotting was performed with primary antibodies overnight at 4 °C. Antibodies included LMNA (Abcam, SAB4200236), LMNB (Abcam, ab16048), *β*-actin (Proteintech, 66009). Quantification was carried out using ImageJ by comparing the intensity of each marker band in a rectangular selection with a fixed size to that of the corresponding reference protein, *β*-actin in the same lane.

### Intracellular optical-tweezers based microrheology

Cells were plated on MatTek glass-bottom dishes coated with matrigel. 2 *µ*m-diameter fluorescent beads (Invitrogen) were added to the cell culture medium at a 1:5000 dilution and were incubated with the cells for 60 hours at 37°C with 5% *CO*_2_. About 1 to 5 beads were internalized depending on the GBM cell line. Cells were stained with Hoechst (33384) at a 1:10000 dilution in cell culture medium for 15 mins before the nuclear rheology experiment.

The setup coupling optical tweezers and fast confocal microscopy has been described in detail previously [34]. In brief, a single fixed optical trap was created by connecting a 1060-1100 nm infrared laser beam (2 W maximum output power; IPG Photonics) to the back port of an inverted Eclipse microscope (Nikon) equipped with a resonant laser confocal A1R scanner (Nikon), a 37 °C incubator, and a nanometric piezostage (Mad City Labs).

The protocol for nuclear indentation experiments and the method for analysis was described in [35]. Briefly, to indent the nucleus, a bead that initially contacted the nucleus was first trapped with the laser. The piezostage was then moved at a constant speed (2.5 *µm* in 1 min) in order to push the nucleus towards the bead. The force was deduced from the displacement of the bead from the trap center (trap stiffness 240 pN/*µ*m) and the indentation depth was obtained by image analysis. A visco-elastic model was used to deduce the nuclear elasticity from the force-indentation curves [35].

### Immunofluorescence and confocal imaging

GBM and RG cells were seeded on matrigel-coated coverslips (33 *µ*g/ml) overnight. Cells were fixed in 4% paraformaldehyde (Thermo Fisher) in PBS, then permeabilized with 0.1% Triton-X100 for 10 min and blocked with 3% BSA-PBS (Sigma) for 1h. Primary antibodies were diluted in 3% BSA-PBS and incubated for 2 hours at room temperature. Secondary antibodies were diluted 1:400 in 3% BSA-PBS and incubated for 1 hour at room temperature. Antibodies used are listed in the Supplementary Methods (Table S1).

Confocal microscopy was performed on a Zeiss LSM780 laser-scanning confocal microscope equipped with white a light laser (WLL), a 405 nm diode laser, three Internal Spectral Detector Channels (PMT), and two Internal Spectral Detector Channels (HyD) GaAsP. Sequential confocal images were acquired using a 20x water-immersion objective (Plan-Apochromat 20x/1.0). For each staining, the acquisition settings (i.e., laser power, beamsplitters, filter settings, pinhole diameters, and scan mode) were kept the same. ImageJ was used to process images (see Supplementary Methods for the definition of the shape parameters extracted by ImageJ).

### 2D in vitro proliferation and migration assay and image analysis

Cells were plated on 6-well plates coated with matrigel at an initial concentration of 4000 cells/cm^2^. The lens-free holographic device Cytonote 6W (Iprasense, Helioparc, France) was used to quantify cell proliferation and to track 2D cell migration. Image acquisition was started 2 hours after cell seeding. Images were recorded every 20 min for 72 hours. The device allows a large recording field of view (29.4 mm^2^). Images were opened as stacks in Fiji and cells were detected automatically using the Trackmate plugin([36]). Three .csv files(“Spots”, “Edges” and “Tracks”) were extracted for each experiment. More details on the extraction of migration parameters are given in Supplementary Methods (Table S2). A custom Matlab code was used to finalize graph visualization and statistical analysis.

### In vivo invasion assay

*In vivo* analysis of GBM cells invading the zebrafish brain was performed as previously described [37, 38]. In brief, zebrafish larvae of Tg(fli1a:rfp) to mark endogenous blood vessels were obtained from fertilized zebrafish eggs 3 days prior to xenotransplantation. Larvae were made transparent by preventing melanin pigmentation using N-phenylthiourea (PTU) (0.003% final). Larvae of 3 dpf were mounted in 2.5 mm wide V-shaped agarose trenches and microinjected after 160 mg/L tricaine treatment using a mechanical micromanipulator (CellTram oil vario microinjector, 5176000.025, Eppendorf) with GBM cells expressing GFP. 2.5 % of a an 80 % confluent 10 cm Petri dish of GBM cells in 5 µl PBS was microinjected into the zebrafish Optic Tectum just above the Middle Cerebral Vein at a maximum of 100 µm from the surface. Xenografts containing a single tumor mass formed by 20-50 cells located in the top 200 µm from the Optic Tectum were selected. After 4 hours of recovery at 32 °C, the larvae were mounted for imaging in a 1% low-melting agarose solution in a 35 mm diameter glass-bottom video-imaging dish. A Nikon Ti2E spinning-disk confocal microscope was used to image the tumor cell mass and invaded cells at 4h and 72h post-injection. The Imaris software was used to segment the cells and calculate the 3D area occupied by the tumor cell mass and invaded cells. The invasion index was calculated as the ratio between the averaged distance travelled by the top 10 cells at 72 hours compared to 4 hours.

### Statistical analysis

All data are presented as mean values ± standard error mean (s.e.m.) of at least three (N≥3) independent experiments. In nuclear rheology experiments, at least n*>*30 nucleus were measured in each condition. Violin plots show the median value (dashed line), and the first and third quartiles (dotted lines). Error bars correspond to s.e.m. Statistical relevance was evaluated using Student’s t-test or one-way ANOVA test with GraphPad Prism, depending on the number of cells, number of samples, and the normality of the distribution. For protein expression levels measured from Western Blot experiments, a one-way ANOVA test was used to compare the protein levels in each GBM cell line to RG cells. For protein downregulation by siRNA, a t-test was used to compare to the siLuciferase control. p-values are reported as non-significant (n.s. p *>* 0.05, p-values are indicated), or significant *p *<* 0.05, **p *<* 0.01, ***p *<* 0.001, ****p *<* 0.0001).

Correlation analysis was performed using Spearman’s rank correlation analysis. Curve-fitting was done using either linear fits or nonlinear fits including one-phase decay with or without plateau. Correlation and curve fitting were performed with GraphPad Prism.

## Results

### Patient-derived glioblastoma cell lines upregulate lamin A

We have previously shown that a grade IV GBM cell line exhibits elevated expression of lamin A and a stiffer nucleus compared to a grade III counterpart [35]. This observation prompted us to study the potential influence of nuclear mechanics on GBM aggressiveness, with a focus on the role of lamin proteins. We selected eleven patient-derived GBM cell lines. These include four proneural subtype cell lines: U3008, U3013, U3021, and U3047; four classical subtype cell lines: U3009, U3017, U3039, and U3123; and three mesenchymal subtype cell lines: U3031, U3065, and U3088. As a non-tumoural control, we used radial glial (RG) cells.

We found that lamin protein expression exhibits distinct patterns between the patient-derived GBM cell lines and RG cells. Specifically, lamin A expression levels are higher in all GBM cell lines, whereas RG cells show very low expression (Fig. 1a-b). Nine out of the eleven cell lines demonstrate a significantly higher lamin A expression levelcompared to RG cells (Fig. 1c). The expression pattern of lamin C closely mirrors that of lamin A (Fig. 1d, Supplementary Fig. S1a, Supplementary Table S3). However, since RG cells display higher expression of lamin C than lamin A, lamin A appears to be a more suitable biomarker to distinguish GBM cells from RG cells.

**Figure 1:**
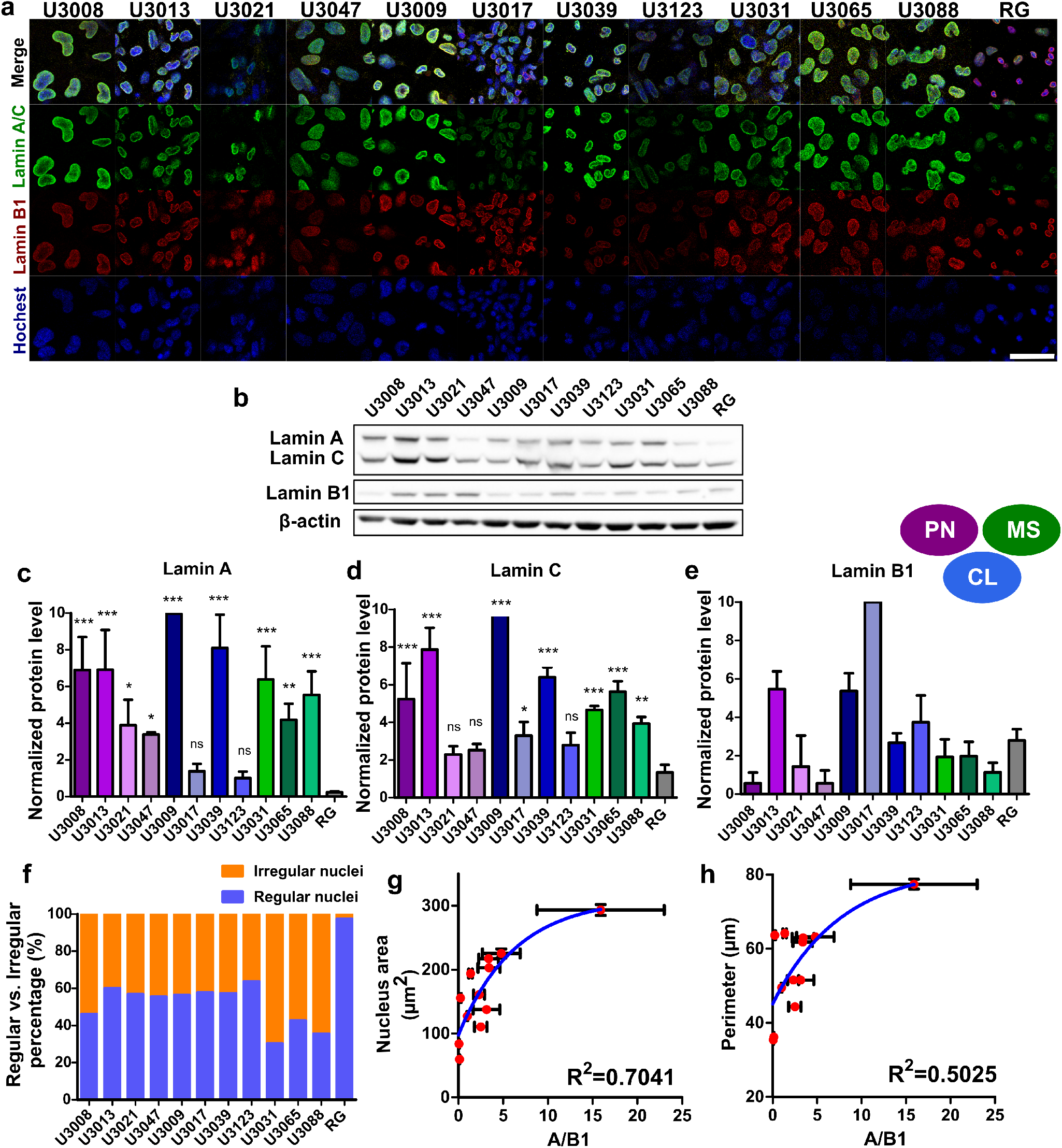
A- and B-type lamins are differently expressed in GBM cells and have opposing effects on nuclear morphology in GBM cells. **a** Immunostaining images of GBM cell lines and RG cells. Lamin A/C (green), Lamin B1 (red). DNA was counterstained with Hoechst (blue). Scale bar 50 *µ*m. **b** Western blot of lamin A/C and lamin B1.*β*-actin was used as a loading control. **c-e** Densitometric analyses for lamin A, lamin C, and lamin B1 respectively. The graphs are shown as mean values ± SEM of three different experiments. Asterisks indicate statistically significant differences with respect to control RG cells; one-way ANOVA test, n.s. p *>* 0.05, *p *<* 0.05, **p *<* 0.01, ***p *<* 0.001. Colors indicate the GBM subtype (purple, proneural PN; blue, classical CL; green, mesenchymal MS). **f** Proportions of nuclei displaying regular vs. irregular shapes in GBM cells and RG cells. At least 100 nuclei were analyzed for each cell line. **g** Correlation between the nuclear area and the lamin ratio A/B1. The fitting curve follows a one-phase decay. Spearman test *R*^2^=0.8322, p=0.0008 (***). **h** Correlation between the nuclear perimeter and the lamin ratio A/B1. The fitting curve follows a one-phase decay. Spearman test R=0.5874, p=0.0446 (*).

The expression of lamin B1 exhibited more variability and did not correlate with lamin A expression (Supplementary Fig. S1b). Some GBM cell lines exhibited increased expression of lamin B1 compared to RG cells, while others showed decreased expression levels (Fig. 1e). Together these results show that, among the three lamin proteins studied, lamin A emerges as the key discriminator between GBM and RG cells.

### A- and B-type lamins have opposing effects on nuclear morphology

By visualizing the nucleus and lamin proteins, we found that nuclear morphology is very het-erogeneous between GBM cell lines and between GBM and RG cells (Fig. 1a). We categorized nuclear shapes into two groups: regular and irregular which encompassed features such as elongated, donut-shape, curved and blebbing nuclei (Supplementary Fig. S1c-d). In GBM cells, we observed that at least 40% of the nuclei display irregular shapes, with some cell lines exhibiting a majority (60–70%) of irregular nuclei (Fig. 1f). Conversely, in RG cells, less than 3% of nuclei are found to be irregular. In addition to exhibiting abnormal shapes, maximum projections of confocal images unveiled significant alterations in nuclear size. Overall, GBM cells display larger nuclear areas compared to RG cells. For instance, the average nucleus projected area of U3008 cells is approximately 300 *µ*m^2^, four times larger than the nuclear size of RG cells.

We then asked whether lamin levels could play a role in the nuclear morphology of GBM cells. We found a weak positive correlation between nuclear size and the expression levels of lamin A and lamin C and a weak negative correlation between nuclear size and the expression levels of lamin B (Supplementary Fig. S2). Interestingly, the size of the nucleus strongly correlates with the ratio between lamin A and lamin B1 (Fig. 1g-h). Nonlinear regression analyses (one-phase decay) revealed a significant correlation between the nuclear projected area or perimeter and the ratio between lamin A and lamin B1 expression levels (termed A/B1 ratio in the following) across the eleven GBM cell lines and RG cells (Fig. 1g,h, Supplementary Tables S4, S5), indicating that the influence of lamin expression on cell nuclear size is not restricted to a specific cell line. We did not observe similar correlations among GBM cells between the expression levels of lamin proteins and the proportion of irregular nuclei (Supplementary Fig. S3) or nuclear morphological parameters, such as circularity, roundness, and shape index (Supplementary Fig. S4). Both increases in nuclear-projected area and perimeter as a function of the A/B1 ratio are well described by an exponential saturation with a plateau value at high (*>* 20) A/B1 ratios:

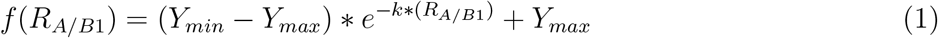

where *R*_*A/B*1_ is the A/B1 ratio, and *Y*_*min*_ and *Y*_*max*_ are the minimal and maximum nuclear-projected area or perimeter, respectively. Curve fitting gives *Y*_*min*_ = 97.59*µm*^2^, *Y*_*max*_ = 311.4*µm*^2^, *k* = 0.1588 for the nuclear projected area and *Y*_*min*_ = 45.14*µm, Y*_*max*_ = 82.09*µm, k* = 0.1293 for the nuclear projected perimeter.

### Lamin A and Lamin B1 increase nuclear elasticity collaboratively

Because GBM cells and RG cells exhibit variable lamin levels which correlate with changes in nuclear morphology, we next asked whether nuclear mechanics also differ in these cell populations. To address this issue, we used optical tweezers (OT) microrheology [34, 35] to quantify the elasticity of the nucleus. Nuclear indentation experiments (Supplementary Movies S1-S2) allowed us to distinguish between ‘soft’ and ‘stiff’ nuclei (Fig.2a,b). Notably, softer nuclei exhibited heightened susceptibility to deformation, resulting in an extended retention time of laser-held beads during nuclear movement (2.5*µ*m/60s). Conversely, stiffer nuclei exhibited reduced compliance, causing early escape of the bead from the laser trap. Consequently, for soft nuclei, we observed pronounced nucleus deformations coupled with small bead displacements from the optical trap center, corresponding to low forces, while stiff nuclei displayed reduced nucleus deformations and large bead displacements from the optical trap center corresponding to large forces (Fig.2a,b). The nucleus elasticity was measured by fitting the force-indentation curves with a visco-elastic model [35].

Our results show that the elasticity of the nucleus varies significantly between distinct cell lines. Specifically, within the cohort of patient-derived GBM cell lines, a comparison between the regular and irregular nuclei subgroups unveiled inherent differences in nucleus elasticity. In the regular nuclei group, GBM cell lines consistently exhibit larger nucleus elasticity than RG cells. The difference is particularly pronounced in U3013, U3009, and U3039 cell lines, for which significant differences with RG cells are observed (Fig. 2c). In the irregular nuclei group, nuclear stiffness is more variable within a given cell line (Fig. 2d), probably because of the local nature of the measurement. This observation underscores the complex interplay between nuclear architecture and cellular heterogeneity in the context of GBM. Remarkably, we found that the elasticity of irregular nuclei is generally lower than that of regular nuclei within the same cell line (Fig. 2e). However, two exceptions were noted in cell lines with low lamin levels, U3017 and U3047.

**Figure 2:**
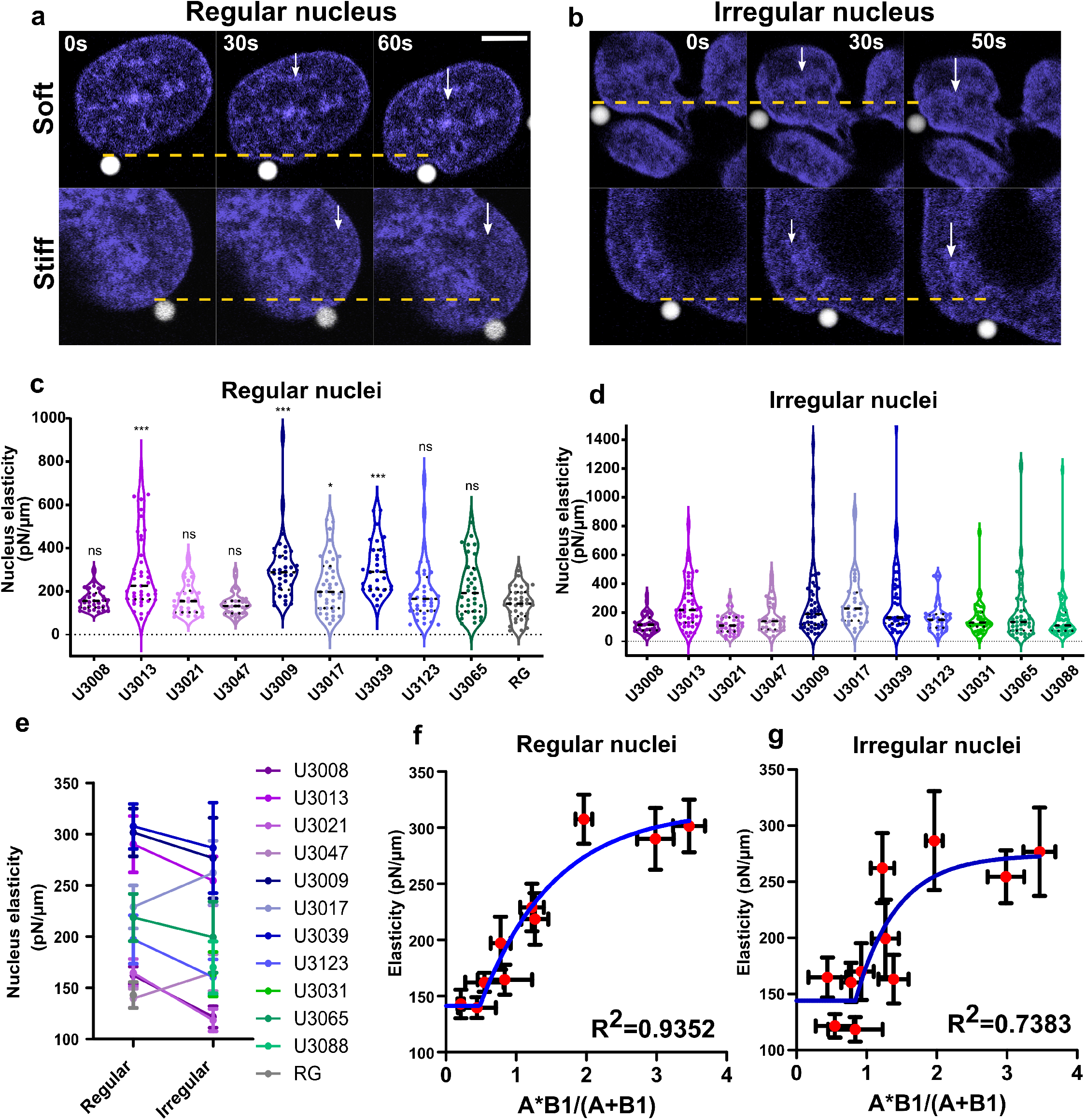
Differences in nuclear elasticity between GBM cell lines correlate with lamin levels. **a**,**b** Time-lapse images of regular (a) and irregular (b) nucleus indentation. The white arrow indicates the direction in which the nucleus in moved. The dotted line marks the initial position of the fluorescent bead in the center of the trap. Bead diameter: 2 *µm*. Scale bar 5 *µ*m. **c** Violin plot of OT measurements of the elasticity of regular nuclei in 9 GBM cell lines and in RG cells (N≥3, n≥30 cells). U3031 and U3088 cells did not have a sufficient number of regular nuclei for measurement. **d** Violin plot of OT measurements of the elasticity of irregular nuclei in 11 GBM cell lines (N≥3, n≥30 cells). RG cells did not have a sufficient number of irregular nuclei for measurement. **e** Comparison of the OT measurement of nuclear elasticity for regular and irregular nuclei in each cell line. **f** Correlation between the elasticity of regular nuclei and the lamin ratio (A*B1)/(A+B1). The fitting curve follows a one-phase decay with an initial plateau. Spearman test R^2^=0.9273, p=0.001 (***). **g** Correlation between the elasticity of irregular nuclei and the lamin ratio (A*B1)/(A+B1). The fitting curve follows a one-phase decay with an initial plateau. Spearman test R^2^=0.7273, p=0.0112 (*).

We found that nuclear elasticity weakly correlates with the levels of lamin A, lamin B1 and lamin C, and the ratio A/B1 (Supplementary Fig. S5a-c), while it correlates with the sum of the expression levels of lamin A and lamin B1 denoted as *A* + *B*1 (Supplementary Fig. S5d, Supplementary Tables S6, S7). Interestingly, we found that nuclear elasticity correlates much more strongly with a lamin level *L* defined by 1*/L* = 1*/A* + 1*/B*1 or *L* = (*A* * *B*1)*/*(*A* + *B*1), where *A* is the expression level of lamin A and *B*1 is the expression level of lamin B1. Assuming that the two ‘layers’ of Lamin A and Lamin B1 act mechanically as springs in series, *L* represents the effective lamin level which may participate in nuclear mechanics. Consistently, when plotted as a function of the effective lamin level *L*, we found that the nuclear elasticity *K* measured by OT for both regular and irregular nuclei increases with *L* (Fig. 2f-g, Supplementary Tables S8, S9) and is well adjusted by a one-phase decay non-linear regression corresponding to:

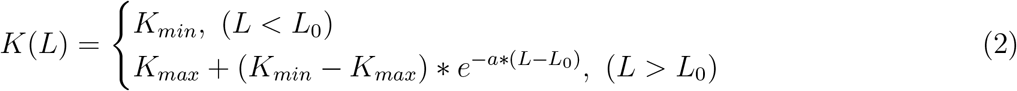

- *K*_*min*_ denotes the minimum elasticity of the nucleus at low *L* values.
- *L*_0_ is the critical effective lamin level above which nuclear elasticity is sensitive to *L*.
- *a* denotes the cell sensitivity to the effective lamin level *L*, representing the rate at which nuclear elasticity increases with *L*.
- *K*_*max*_ is the maximum elasticity achievable by the nucleus at high lamin levels. Once the nucleus reaches this value, further increase in lamin levels does not result in significant increase in elasticity.

For regular nuclei, the fit parameters are *L*_0_ = 0.48, *a* = 0.95, *K*_*min*_ = 141.4, and *K*_*max*_ = 316.5 and nuclear elasticity ranges from 140 to 307 *pN/µm*, from U3047 to U3009 GBM cell lines, respectively. The effective lamin level *L* exhibits a range of 0.2 to 3.5, from RG cells to the U3009 GBM cell line.

For irregular nuclei, the fit parameters are *L*_0_ = 0.84, *a* = 1.69, *K*_*min*_ = 144.1, and *K*_*max*_ = 274.1 and nuclear elasticity ranges from 119 to 287 *pN/µm*, from U3021 to U3039 GBM cell lines, respectively. The effective lamin level *L* exhibits a range of 0.5 to 3.5, from U3008 to U3009 GBM cell lines, respectively.

Regarding the minimum stiffness of the nucleus *K*_*min*_, our analysis indicates a similar basal elasticity for both regular and irregular nuclei. Regarding *L*_0_, in the regular group, nuclear elasticity initiates its increase when the effective lamin level reaches 0.48. In contrast, within the irregular group, the effective lamin level needs to reach 0.84 before influencing nuclear stiffness. Regarding *a*, interestingly, nuclear elasticity increases faster with *L* in the irregular group compared to the regular group. Regarding *K*_*max*_, we found that the plateau value of irregular nuclei is 10% smaller than that of regular nuclei, suggesting a lower maximum elasticity threshold in irregular nuclei compared to their regular counterparts.

### GBM cell proliferation correlates positively with Lamin A expression levels

Our results so far show that lamin expression levels in patient-derived GBM cell lines are associated with nuclear morphology and elasticity. In contrast with other cancer types for which tumour progression is characterized by cell and nucleus softening [12, 29], we have previously reported that a grade IV GBM cell line exhibits a stiffer nucleus than a grade III glioma cell line [35]. Hence, we aimed here to investigate in more detail the impact of lamin A on GBM aggressiveness in patient-derived cell lines.

We first measured cell proliferation rates, as an indicator of aggressiveness, using lens-free microscopy in the eleven patient-derived GBM cell lines and in RG cells (Supplementary Movie S3) and found very different proliferation profiles (Fig. 3a-b). For instance, U3088 cells, U3008 cells, and RG cells show high, medium, and low proliferation rates, respectively. Cell lines with low proliferation rates exhibit a 1.5-2 fold increase in cell number in 72 hours, while cell lines with high proliferation rates reach up to a 3-5 fold increase in cell number (Fig. 3b). Notably, RGs cells proliferate the slowest among the twelve cell lines we studied.

**Figure 3:**
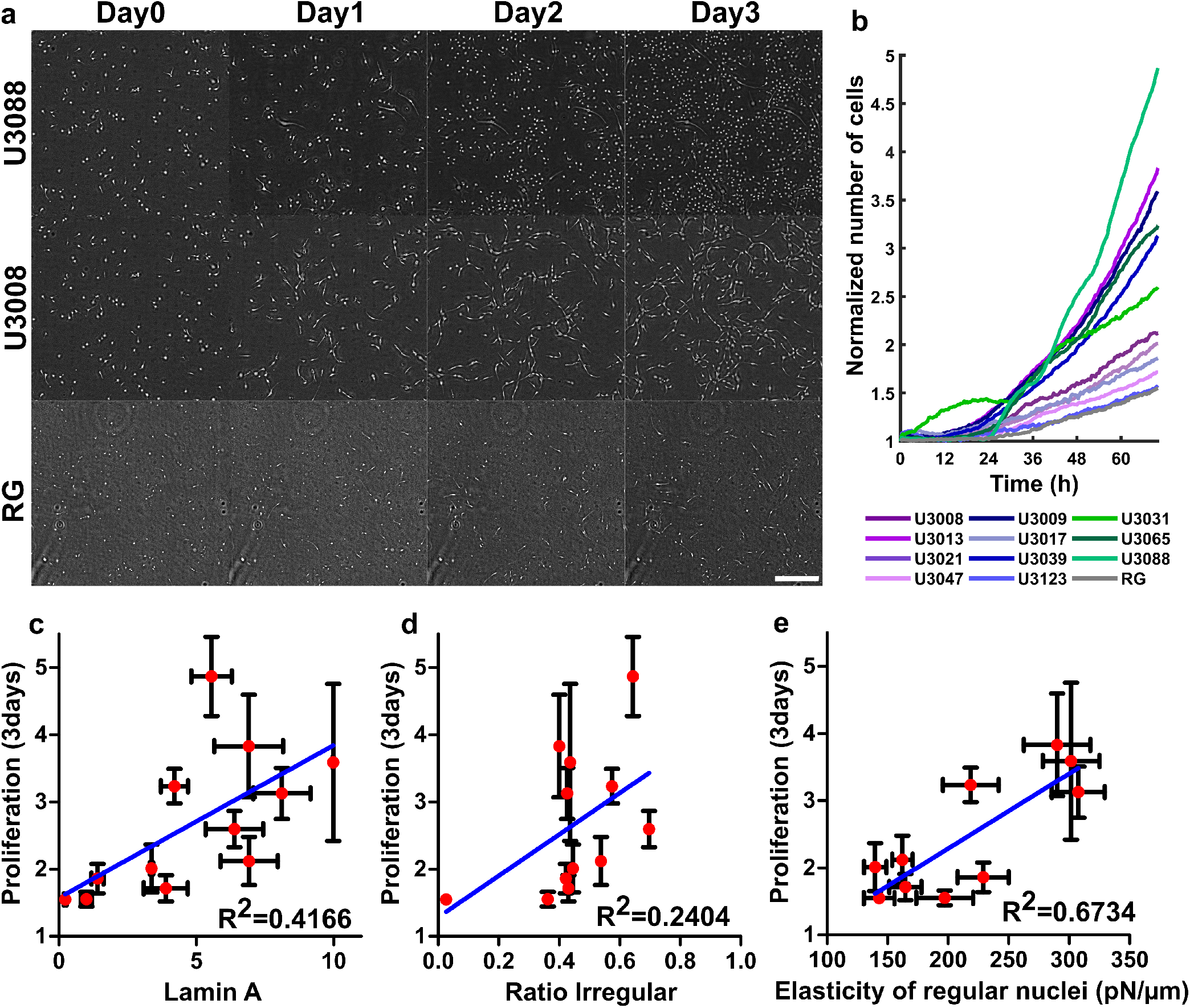
GBM cell lines expressing high levels of lamin A show elevated proliferation rates. **a** Brightfield lens-free microscopy images of 2D cell cultures followed for three days. U3088, U3008, and RG cells represent high, medium, and low proliferation rates, respectively. Scale bar 500 *µ*m. **b** Time evolution of cell proliferation on a 2D substrate during three days. (N=3 independent experiments, n≥2000 cells). For a better readability of the figure, error bars are not shown but can be appreciated from panels c-e.**c** Correlation between proliferation and lamin A levels. **d** Correlation between proliferation and the proportion of irregular nuclei. **e** Correlation between proliferation and nucleus elasticity.

We then asked whether lamin expression levels correlate with the proliferation rate (Figure 3c, Supplementary Fig. S6a). Plotting the proliferation rate against the lamin A expression level revealed a statistically significant positive correlation (Supplementary Table S10): cell lines with higher lamin A expression consistently exhibit higher proliferation rates, while those with low lamin A expression display lower proliferation rates. Figure 3c illustrates that RG cells and the four GBM cell lines with low lamin A expression (below 4 a.u.) exhibit less than a 2-fold increase in cell number in 72 hours, while the remaining seven cell lines, with lamin expression ranging from 4 a.u. to 10 a.u., exhibit a 2 to 5-fold increase in cell number. A similar correlation was observed with lamin C expression levels (Supplementary Table S11) but not with lamin B1 expression levels (Supplementary Fig. S6a) or with the combinations of lamin levels (Supplementary Fig. S6c)

Given our results showing a link between lamin levels and nuclear morphology and mechanics (Figs. 1 and 2, Supplementary Figures S1-S5), we expected that nuclear morphological and mechanical characteristics could impact on cell proliferation. We indeed found that the proportion of irregular nuclei positively correlates with the rate of cell proliferation, as depicted in Figure 3d, although not in a stastically significant manner (Supplementary Table S12). Similarly and not surprisingly since nuclear elasticity and lamin A expression levels strongly correlates (Supplementary Fig. S5a-b), proliferation rates positively correlate with the elasticity of regular nuclei (Figure 3e, Supplementary Table S13) and, although to a lesser extent, of irregular nuclei (Supplementary Fig. S6b, Supplementary Table S14).

### 2D migration velocity of GBM cells correlates positively with Lamin A expression levels

GBM clinically display diverse motile and invasive capacities. Here, we first used migration on 2D substrates as a second indicator for the aggressiveness of our patient-derived GBM cell lines. The 2D migration of single cells was tracked for three days for the eleven GBM cell lines and for RG cells using lens-free microscopy. RG cells are the slowest moving cells with an average speed of 0.3 *µ*m/min and GBM cell lines migrate much faster and exhibit a range of migration velocities (Fig 4a). 2D migration velocity increases with increasing lamin expression levels (Fig 4b, Supplementary Fig. S7a). With the exception of two GBM cell lines of the classical subtype with low lamin A expression levels, 2D migration velocity strongly correlates with lamin A expression levels (Fig 4b). We found that 2D cell migration velocity weakly correlates with the expression levels of lamin B1 and lamin C or the combinations of lamin levels (Supplementary Fig. S7a-b). Interestingly, we found that the 2D migration velocity correlates linearly with nuclear elasticity (Fig. 4c, Supplementary Tables S15, S16).

**Figure 4:**
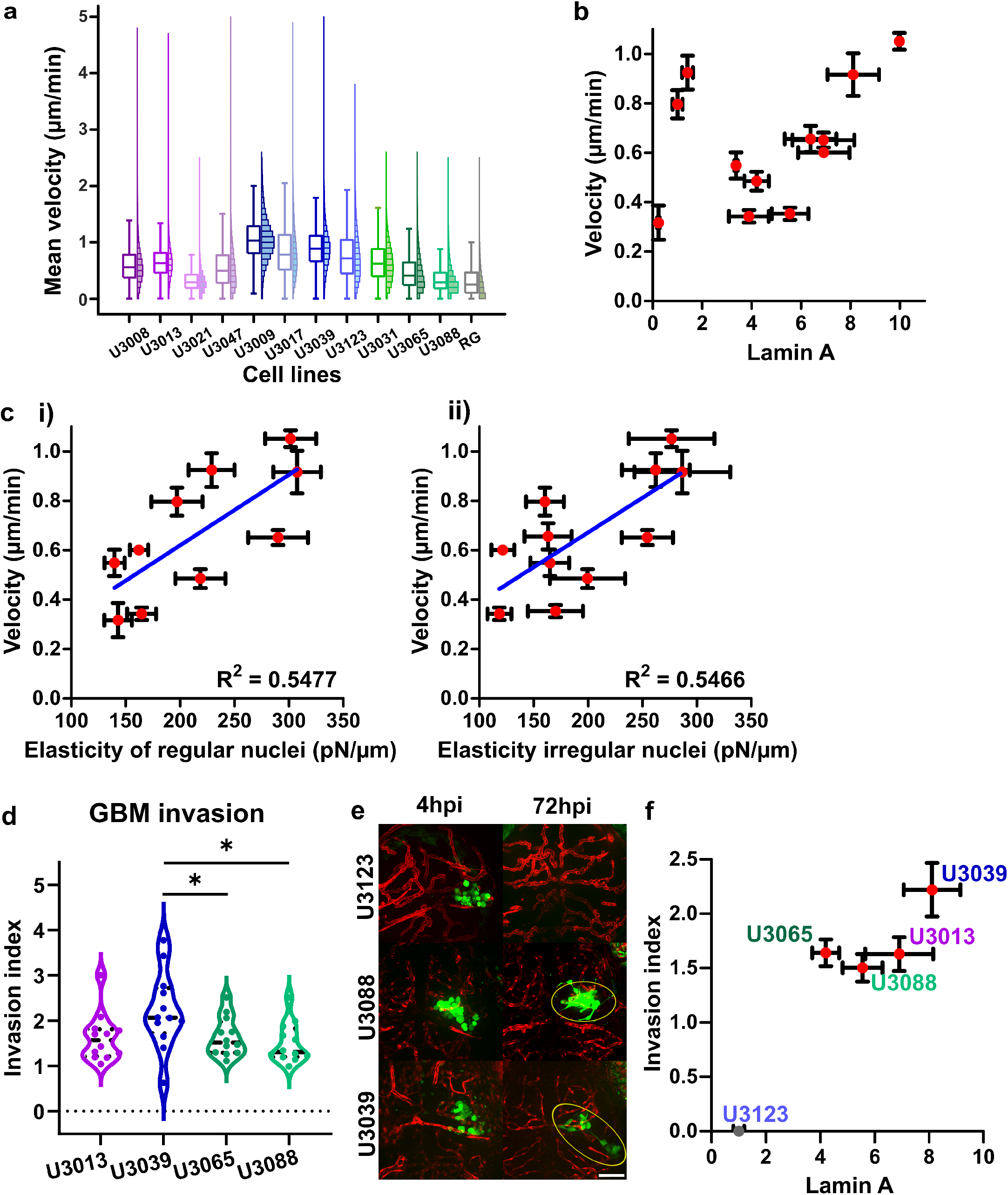
GBM cell lines expressing high levels of lamin A show elevated 2D migration velocity and higher invasion in the zebrafish brain. **a** Histo-box plot of cell migration velocity. The results show the mean velocity along the track of single cells during 3 days (N=3 independent experiments, n≥2000 cells). **b** Correlation between migration velocity and lamin A level. **c** Correlation between cell velocity and nuclear elasticity for regular nuclei (i) and for irregular nuclei (ii). **d** Violin plot of the invasion index of four GBM cell lines (U3013, U3039, U3065, U3088) in the zebrafish brain. **e** Fluorescence images of GBM cell lines (U3123, U3088, U3039) invasion in the zebrafish brain (GBM cells in green, endothelial cells delineating blood vessels in red) at 4 hours and 72 hours post-injection (hpi) (N=8 zebrafish larvae, n≥10 cells). Yellow circles in the 72hpi U3088 and U3039 images indicate the zone of invasion. U3123 cells do not survive in the zebrafish brain. Scale bar, 50 µm. **f** Correlation between the expression level of lamin A and the invasion index in the zebrafish brain of the four GBM cell lines shown in d). Note that we added the U3123 data with a zero invasion index, as this cell line does not survive in the zebrafish brain.

*In vivo*, the proliferation rate and migration velocity are two important indicators of the degree of malignancy of a tumour, as they determine the growth rate and metastatic potential of a tumour. These two indicators are not necessarily related. Indeed, when the data of the three GBM subtypes were pooled, we did not observe any clear correlation between proliferation rate and 2D migration velocity (Supplementary Fig. S7c). However, once grouped by subtype, 2D cell migration velocity and proliferation rate positively correlate for cell lines of the proneural and classical subtypes, while they were negatively correlated for cell lines of the mesenchymal subtype (Supplementary Fig. S7c). Interestingly, cells of the classical subtype move twice as fast as those of the proneural subtype (Supplementary Fig. S7c).

### Impact of lamin A on glioblastoma invasion

Our results show that lamin A levels are associated with both GBM cell proliferation and 2D migration (Figs. 3 and 4a-c). However, proliferation and migration on a 2D stiff substrate is not always indicative of the invasion rate in three-dimensional tissues. We thus asked whether lamin A expression levels could affect the invasiveness of GBM cells. To address this question, we injected cancer cells into the brain of zebrafish and compares the invasion of five different GBM cell lines. These cell lines (U3013, U3039, U3065, U3088, and U3123) were selected for their diversity in their expression levels of lamin A, the mechanical and morphological properties of their nucleus, and their behaviours in the 2D cell proliferation and migration assays. We found that these five GBM cell lines had a very different invasion rate (Fig.4 d) as quantified using their invasion index (see Methods). Unexpectedly, the U3123 cell line, which expressed the lowest amount of lamin A, did not invade the zebrafish brain and died 24 hours after injection (Fig. 4e, d). Furthermore, we observed a positive correlation between the lamin A expression levels and the invasion index for the remaining four cell lines (Fig. 4f). The U3039 cell line, which expresses the highest level of lamin A, had the highest invasion index which was statistically significantly higher than for the U3065 and U3088 cell lines (Fig.4 d, e). Taken together our results demonstrate that, in GBM, the lamin A expression level impacts on nuclear morphology and mechanical properties and positively correlates with cell proliferation and migration in 2D and with cell invasion in the zebrafish brain, which can be considered as indicators of GBM aggressiveness.

### Downregulating lamin A expression reduces nuclear size, nuclear elasticity, proliferation and 2D migration velocity in GBM cells

To further test the potential causal links between lamin levels and GBM aggressiveness, we studied the effects of lamin A downregulation on nuclear morphology and mechanics and on cell proliferation and migration. We used siRNA-mediated downregulation of lamin A (si-Lamin A) in two GBM cell lines, U3009 and U3039 which are both of the classical subtype and have the highest lamin A expression levels among our eleven GBM cell lines. As a control, siRNA against Luciferase (si-Luciferase) was used. We checked the effect of lamin A downregulation by immunostaining and Western blot (Fig. 5a, b). Quantification of the Western blot results show that the lamin A expression levels was decreased by 30-40% compared to the control group in both cell lines three days after siRNA transfection (Fig. 5c).

**Figure 5:**
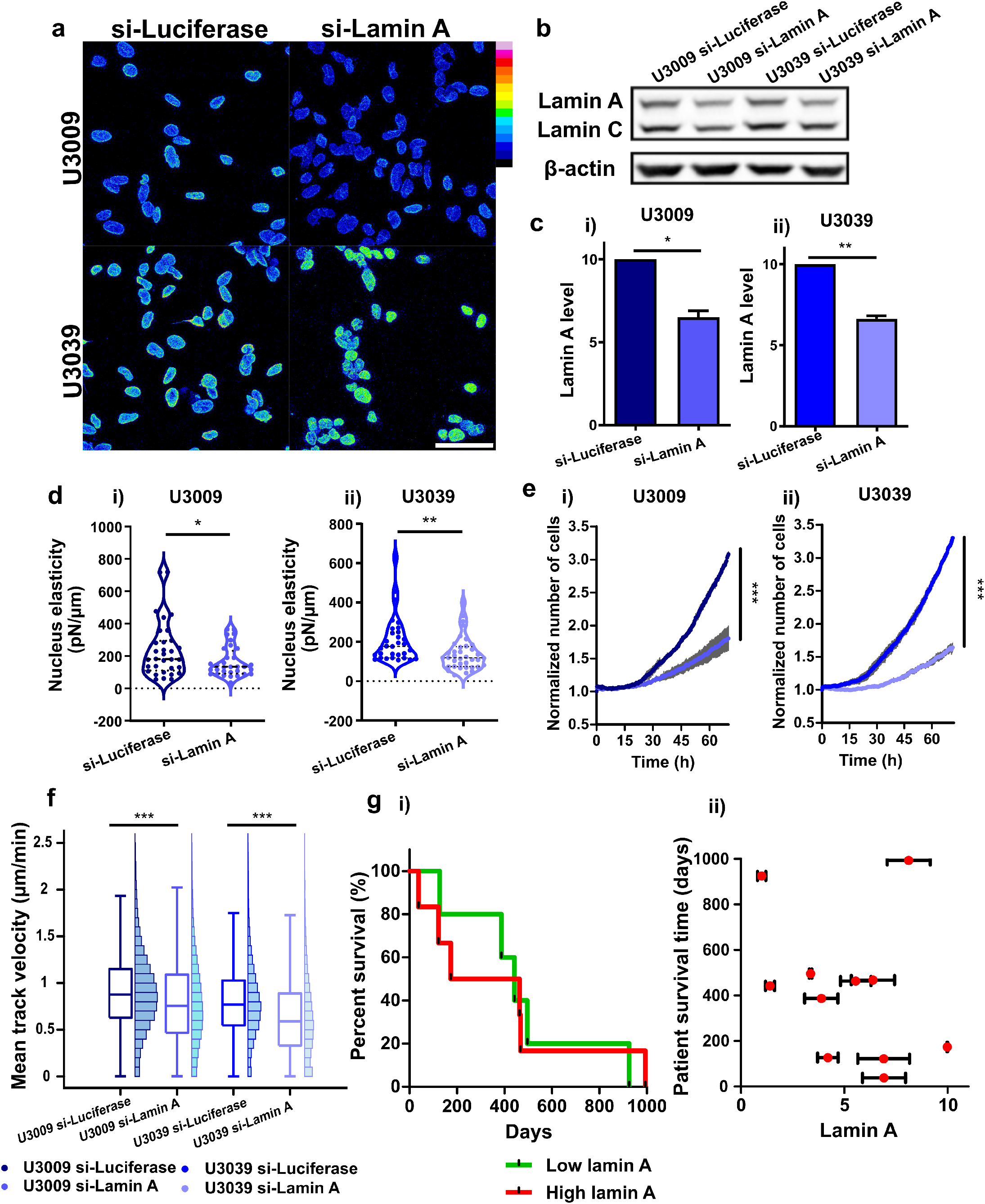
Downregulation of lamin A induces a decrease in nuclear elasticity, proliferation and 2D migration. **a** Immunostaining images of U3009 and U3039 cells treated with si-Lamin A and si-Luciferase as a control. Scale bar, 50 *µ*m. **b** Western blot showing the reduction of lamin A expression in si-Lamin A treated cells compared to control si-Luciferase treated cells. *β*-actin was used as a loading control. **c** Quantification of lamin A levels from the Western blot results. Both U3009 (i) and U3039 (ii) cells treated with si-Lamin A show a 30-40% decrease in lamin A expression compared to control cells. **d** Violin plots of nuclear elasticity measured by intracellular OT for U3009 (i) and U3039 (ii) cells treated with si-Lamin A or control si-Luciferase (N≥3, n≥30 cells). **e** Time evolution of cell proliferation on 2D substrates during three days for U3009 (i) and U3039 (ii) cells treated with si-Lamin A or control si-Luciferase (N = 3, n≥2000 cells). **f** Histo-box plot of cell migration velocity for U3009 and U3039 cells treated with si-Lamin A or control si-Luciferase. The results show the mean velocity along the track of single cells during three days (N = 3, n≥2000 cells). **g** Clinical data of patient survival. i) Kaplan-Meier plot of patient survival percentage for GBM patients expressing high levels of lamin A (U3088, U3031, U3013, U3008, U3039, U3009) or low levels of lamin A (U3123, U3017, U3047, U3021, U3065). ii) Correlation between patient survival time (in days) and lamin A expression levels. In panels (a-f), the graphs show mean values ± SEM of three different experiments. Asterisks indicate statistically significant differences between si-Lamin A and si-Luciferase treated U3009 or U3039 cells; t-test, n.s. p > 0.05, *p < 0.05, **p < 0.01, ***p < 0.001.

**Figure 6:**
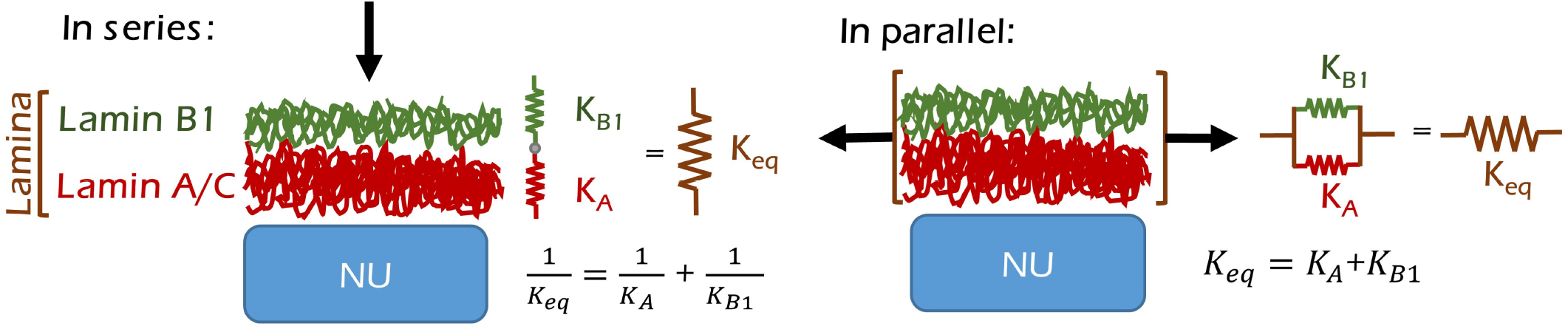
Mechanical models of the nuclear lamina. The two layers of lamin A and lamin B1 can act mechanically as springs in series (left panel) or in parallel (right panel).

We first investigated the effects of lamin A downregulation on nuclear morphology and found that both U3009 and U3039 cells treated with si-Lamin A have smaller nuclei than control cells (Supplementary Fig. S8). In the case of U3039 cells, the nucleus projected area is reduced by 30% after lamin A downregulation. In the case of U3009 cells, lamin A downregulation does not induce any change in the nucleus projected area but decreases the nucleus thickness by around 30% (Supplementary Fig. S8).

We next asked whether the mechanical properties of the nucleus are affected by lamin A downregulation. We observed a 30% decrease in nuclear elasticity following lamin downregulation (Fig. 5d), showing that lamin A contributes to nuclear elasticity and explaining the correlation between lamin A levels and nuclear elasticity (Supplementary Fig. S5).

We also measured the impact of lamin A downregulation on two indicators of GBM aggressiveness, 2D cell proliferation and migration velocity. We found a strong decrease in the proliferation rate of both U3009 and U3009 cells treated with si-Lamin A compared with the control (Fig. 5e). Note that, even with a modest (30%) decrease in lamin A expression (Fig. 5a-c), the proliferation rate is decreased to the same level as RG cells, our control cells. Similarly, we measured a significant decrease in 2D migration velocity after lamin A downregulation (Fig. 5f). These results demonstrate that decreasing lamin A expression levels can efficiently reduce GBM cell proliferation and migration in 2D.

Finally, to test the potential clinical relevance of our results, we correlated the lamin A expression levels of the primary cell lines with the survival time of the corresponding patients. Although the survival time strongly depends on the age at which the patient is first diagnosed with GBM, the survival of patients with low lamin A levels (U3123, U3017, U3047, U3021, U3065) was higher than that of patients with high lamin A levels (U3088, U3031, U3013, U3008, U3039, U3009) within the first year after diagnosis (Fig. 5g panel i), although not in a statistically significant manner. Consistently, the patient survival time appears to decrease with the lamin A expression level in ten out of the eleven patient-derived GBM cell lines (Fig. 5g panel ii), with the notable exception of the outlier U3039 cell line. These observations suggest that lamin A expression may be used as a prognosis marker for GBM.

## Discussion

The mechanical properties of the cell and in particular those of its nucleus are thought to be critical for cancer cell migration and invasion. While tumour progression has been associated with cell and nuclear softening in several types of cancers, we have previously shown that, for glioma tumours, a grade IV GBM cell line exhibits a stiffer cytoplasm and nucleus than a grade III cell line [35]. These results lead us to hypothesize that nuclear stiffening may be associated to glioma aggressiveness. Here we test this hypothesis in the context of GBM using eleven patient-derived cell lines and non tumoral RG cells. We find that the regulation of nuclear morphology and stiffness by lamins is associated with GBM cell proliferation, migration, and invasion.

### Lamins and nuclear morphology in GBM cells

Lamin proteins are known to regulate nuclear shape in several cell types. For instance, lamin B deletion has been shown to induce nuclear blebbing [39] in HeLa cells. It was also observed that lamin B1 stabilizes the nucleus shape by restricting outward protrusions of the lamin A/C network [40]. High-resolution imaging has shown that nuclear blebs localize at zones lacking lamin B [40], suggesting distinct roles of lamin A/C and lamin B in nuclear morphology. Lamin A is supposed to expand the nucleus, whereas lamin B contracts it [40, 41]. Previous studies have demonstrated the opposing roles of lamin A and lamin B in determining cell nuclear morphology [42]. Here we confirm these results in GBM cells. We show that the ratio between the expression levels of lamin A and lamin B1 (A/B1 ratio) positively correlates with nuclear size (Fig. 1g,h, Supplementary Fig. S2) and that reducing lamin A expression levels by about 30-40% decreases nuclear size in two GBM cell lines with high lamin A expression levels (Supplementary Fig. S8). We also distinguished between cells exhibiting a ‘regular’ and an ‘irregular’ nucleus (Supplementary Fig. S1c,d) and found that all GBM cells have a much higher proportion of irregular nuclei than RG cells (Fig. 1f) but no correlation was observed with the expression levels of lamin proteins (Supplementary Fig. S3).

### Lamins and the mechanical properties of the nucleus in GBM cells

The role of lamins in the control of nuclear mechanics is well established, in particular in the case of lamin A [43, 44]. In agreement with previous results, downregulating lamin A expression by siRNA in two GBM cell lines expressing high levels of lamin A (U3009 and U3039) reduces nucleus elasticity (Fig. 5d). The contribution of lamin B1 to the stiffness of the nucleus is more controversial but lamin B1 does seem to significantly affect the mechanical properties of the nucleus[45]. Tissue and cell stiffness have been shown to correlate primarily with lamin A expression levels [30]. Furthermore, it was recently shown that the loss of A-or B-type lamins significantly softens the nucleus [42].

Here we show that, the mechanical properties of the nucleus (Fig. 2) depend on both lamin A and lamin B1, suggesting a physical link between the two networks of intermediate filament proteins. Although the lamin A and lamin B1 networks are known to localize in close proximity at the nuclear lamina [40], no direct interaction between lamin A and lamin B1 has been reported so far [27, 46]. From a mechanical point of view, depending on their spatial arrangement, the two lamin networks can act as mechanical viscoelastic modules in series or in parallel. Considering only the elasticity of the two networks, the equivalent elasticity *K* is given by *K* = *K*_*A*_ + *K*_*B*1_ if lamin A and lamin B1 act in parallel and by *K* = *K*_*A*_ * *K*_*B*1_*/*(*K*_*A*_ + *K*_*B*1_), where *K*_*A*_ is the elasticity of the lamin A network and *K*_*B*1_ is the elasticity of the lamin B1 network (Fig.6).

When studying the mechanics of our different GBM cell lines and RG cells, we found that nuclear elasticity increases with the quantity *L* defined by *L* = *A* * *B*1*/*(*A* + *B*1) where *A* is the expression level of lamin A and *B*1 is the expression level of lamin B1 (Fig.2f,g). Nuclear elasticity correlates more strongly with the quantity *L* than with individual lamin levels (*A, C* or *B*1), with the *A/B*1 ratio or with the *A* + *B*1 sum which would correspond to a model in which the lamin networks act in parallel (Supplementary Fig. S5). Our data thus strongly favors a physical model in which the lamin A and lamin B1 networks act as springs in series. This finding is supported by the localization of the two networks as two superimposed layers when imaged with super-resolution microscopy [40], with the lamin B1 localizing more externally than the lamin A network in the nuclear lamina. Such a two-layer spatial arrangement of the two networks is consistent with the idea that B-type lamins tend to contract the nucleus while A-type lamins tend to increase nuclear size [40, 41] as discussed above. The two-layer model of the lamin A and lamin B1 networks may also explain the mechanical coupling between the two networks: by compressing the nucleus, an increase in lamin B1 levels in the outer lamina could induce an increase in lamin A concentration in the internal lamina and a subsequent increase in nuclear stiffness. Note that the precise relative localization of the lamin B1 and lamin A networks is still debated and the two networks may actually be intertwined.

Together our measurements of nuclear elasticity with optical tweezers in living cells point to a synergy between the lamin A and lamin B1 networks in the control of nuclear mechanics. To better understand this synergy, we used a one-phase decay non-linear regression to fit the evolution of the nuclear elasticity *K*(*L*) as a function of the effective lamin level *L* (Equation 2). For both regular and irregular nuclei, we found that above a threshold value *L*_0_, the nuclear elasticity increases from a minimum value *K*_*min*_ to a maximum plateau value *K*_*max*_ with a sensitivity *a* to the effective lamin level *L*. As discussed below, the values of these four parameters (*L*_0_, *K*_*min*_, *K*_*max*_, and *a*) differ between regular and irregular nuclei.

At low lamin levels, below the *L*_0_ threshold, the elasticity of the nucleus is small (*K* ≃ *K*_*min*_) and does not vary significantly with *L*. This observation suggests that our optical tweezers techniques probes mostly the mechanics of a lamin-independent component of the nucleus, probably intra-nuclear material such as chromatin. The value of the minimal elasticity *K*_*min*_ is not only similar for RG cells and GBM expressing low lamin levels but also for both regular and irregular nuclei in these cell lines (Fig. 2f,g and Supplementary Tables S8, S9), strengthening our hypothesis that lamins do not contribute to nuclear elasticity below an effective lamin level threshold *L*_0_. Above the *L*_0_ threshold, the nuclear elasticity increases with the effective lamin level *L*, confirming the known impact of lamins on nuclear mechanics [43, 44, 42]. At high lamin expression levels (*L >> L*_0_, the nuclear elasticity saturates. Interestingly, irregular nuclei exhibit a sharper increase in elasticity with a higher threshold value *L*_0_ and a higher sensitivity *a* and reach a lower *K*_*max*_ value (about 10% lower) than than regular nuclei. It has been reported that the lamin B1 network is disorganized or even lacking from regions of local nuclear deformations [40, 47] such as in blebbing, elongated, protruding or donut shapes typical of irregular nuclei (Supplementary Fig. S1c-d). Because our optical tweezers technique probes nuclear mechanics locally, the lower average elasticity we measured for irregular nuclei for most GBM cell lines (Fig. 2c-e) could thus result from heterogeneities in the distribution of the lamin B1 network.

### Lamins and GBM cell proliferation, migration and invasion

Our results not only confirm and specify the role of lamin proteins in the control of nuclear shape and mechanical properties but also highlight their links with other cellular functions in GBM cells. In particular, we demonstrate that the expression level of lamin A correlates with GBM cell proliferation (Fig. 3) and that downregulating lamin A reduces cell proliferation (Fig.5e). Both A-type lamins and B-type lamins have been linked with cell proliferation in previous studies [48, 49, 24]. Silencing lamin A/C in human fibroblasts [48] or knocking-out lamin A in mouse [49] reduces cell proliferation, in agreement with our results. Lamins are known to anchor chromatin at the nuclear envelope and to influence higher-order chromatin organization due to their capacity to bind DNA, chromosomes, and chromatin[50]. Lamins may play a role in the cell cycle-dependent dynamics of nuclear structure since they reversibly separate from chromosomes during mitosis in a phosphorylation-dependent manner [51]. Although the precise mechanisms by which lamin A may promote GBM cell proliferation have to be investigated in more details, our data show a clear link between lamin levels and GBM aggressiveness through cell proliferation.

Another hallmark of GBM cells is their extremely high migratory and invasive capacities [52]. In particular, we have shown previously that cytoplasmic intermediate filaments, in particular vimentin intermediate filaments, play a crucial role in GBM cell migration [52, 38]. Here, we find that 2D migration velocity positively correlates with lamin A levels (Fig. 4b and Fig. 5f). Similarly GBM cell lines which express more lamin A invade more efficiently the zebrafish brain (Fig.4d-f). Because we showed that GBM cell lines with higher lamin levels display a stiffer nucleus, our data suggest that in GBM, in contrast with previous results in other types of cancers such as breast cancer[53, 54], a stiffer nucleus allows more efficient cell migration and invasion. These results are consistent with our previous report that a less aggressive astrocytoma grade III cell line display a softer nucleus than a more aggressive GBM grade IV cell line [35]. Still they may seem surprising when compared to other tumours for which a softer nucleus may allow cells to be more deformable and hence migrate more efficiently through entangled and confined extracellular matrices (ECM) [35, 55, 31]. Our data favor a model in which a stiff nucleus allows cells to push more efficiently through the soft environment of the brain [56].

*In vivo*, GBM cell migration occurs as chain migration along blood vessels, known as vessel co-option[57]. It was recently shown that GBM cells that use vessel co-option to invade the mouse brain exhibit high levels of lamin A [58]. In agreement, in our zebrafish model, we observed that higher lamin A expression correlates with more efficient invasion. This suggests that during chain migration along blood vessels, leader cells with a stiff nucleus, high contractility, and a high matrix degradation activity [38] may ‘dig’ more efficiently through the brain ECM and allow follower cells to migrate in their tracks. Taken together, our data suggest that GBM cells expressing higher levels of lamins show a more aggressive phenotype which may be, at least in part, attributed to an increase in the stiffness of the nucleus.

### Clinical perspectives

In a vast majority of cancer types, including prostate cancer[59], colon cancer[60], gastrointestinal neoplasms[61], gastric carcinoma[62], breast cancer[63] and neuroblastoma[64], decreased lamin A levels have been associated to a higher degree of malignancy[65]. In some cases, the contribution of lamin A to cancer cell invasion is more debated[66]. In contrast, we demonstrate here that lamin levels positively correlate with GBM cell proliferation, migration and invasion, suggesting that up-regulation of lamins, and in particular lamin A, may favor GBM aggressiveness. This specificity of GBM may be explained by the low expression levels of lamins in the healthy brain as compared to other tissues. In organs that are inherently subject to mechanical stress, including the breast, the prostate, the intestine and the pancreas, lamin A is highly expressed in the healthy tissues [30, 29], and decreased lamin A levels is a hallmark of cancer in the tumoral tissues[29]. In the brain, however, lamin A is expressed at very low levels in most cell types including neurons, glial progenitor cells, and astrocytes[67, 68]. Depending on the role and the level of expression of lamins in the tissue of origin, tumour development may thus be characterized by an down-regulation or an up-regulation of lamins. This hypothesis is further supported in the case of the brain by the observation that most LMNA-negative laminopathy patients do not exhibit significant abnormalities or brain developmental defects during fetal development[69], but will later develop cardiac and/or skeletal myopathy. Conversely, for B-type lamins which are strongly expressed in brain tissues compared to lamin A[67, 68], LMNB-negative laminopathy can cause fetal death as well as major defects in the patient’s brain and nervous system during development[70].

Finally, we noticed that survival was higher for patients with low lamin A expression levels one year after diagnosis, and that, with one exception out of the eleven patient-derived GBM cells we studied, the patient survival time decreased with increasing lamin A expression levels (Fig. 5g). Together with our *in vitro* data and our results in the zebrafish brain, this observation further suggests that lamins may serve as clinically relevant therapeutic targets. To conclude, our study highlights the role of lamins in GBM aggressiveness through their impact on nuclear shape and mechanics. Further work should address the potential mechanisms linking the roles of lamins in cell proliferation and cell invasion with their roles in nuclear mechanics, in the context not only of GBM but also of other types of cancers or other diseases.

## Acknowledgments

This work was supported by grants from ITMO Cancer Inserm-Aviesan “Approches interdisci-plinaires des processus oncogéniques et perspectives thérapeutiques : Apports à l’oncologie de la physique, de la chimie et des sciences de l’ingénieur” Edition 2022” (NUTMEG project, grant number 22CP073-00) and from the Labex Who Am I? (ANR-11-LABX-0071) and the “Initiatives d’excellence” (Idex ANR-11-IDEX-0005-02) transverse project BioMechanOE (TP5).

## Author contribution statements

[Jean-Baptiste Manneville] conceptualized the study, and supervised the overall project. [Xiuyu Wang] designed and performed the experiments; collected and analyzed the data; and interpreted the results. [David Pereira] performed optical tweezers experiments. [Isabelle Perfettini, Florent Peglion, Sandrine Etienne-Manneville] provided the zebrafish model and performed invasion experiments in zebrafish. [Ryszard Wimmers, Alexandre Baffet] collected and differentiated radial glial cells. [Ananya Roy, Karin Forsberg-Nilsson] provided GBM cell lines and expertise in GBM biology. [Xiuyu Wang] was responsible for writing the first draft of the manuscript. All authors contributed to the review and editing of the manuscript, approved the final version, and agreed to be accountable for all aspects of the work.

## References

[1] Farina Hanif et al. “Glioblastoma multiforme: a review of its epidemiology and pathogenesis through clinical presentation and treatment”. In: Asian Pacific journal of cancer prevention: APJCP 18.1 (2017), p. 3.

[2] James D Brierley, Mary K Gospodarowicz, and Christian Wittekind. TNM classification of malignant tumours. John Wiley & Sons, 2017.

[3] R Stupp et al. “High-grade glioma: ESMO Clinical Practice Guidelines for diagnosis, treatment and follow-up”. In: Annals of oncology 25 (2014), pp. iii93–iii101.

[4] Sheila K Singh et al. “Identification of human brain tumour initiating cells”. In: nature 432.7015 (2004), pp. 396–401.

[5] Chul-Kee Park, Jeong Mo Bae, and Sung-Hye Park. “Long-term survivors of glioblastoma are a unique group of patients lacking universal characteristic features”. In: Neuro-Oncology Advances 2.1 (2020), vdz056.

[6] Wojciech Szopa et al. “Diagnostic and therapeutic biomarkers in glioblastoma: current status and future perspectives”. In: BioMed research international 2017 (2017).

[7] Kaja Urbańska et al. “Glioblastoma multiforme–an overview”. In: Contemporary Oncology/Współczesna Onkologia 18.5 (2014), pp. 307–312.

[8] Aneta Wlodarczyk et al. “Gaps and doubts in search to recognize glioblastoma cellular origin and tumor initiating cells”. In: Journal of oncology 2020.1 (2020), p. 6783627.

[9] Elena Verdugo, Iker Puerto, and Miguel Ángel Medina. “An update on the molecular biology of glioblastoma, with clinical implications and progress in its treatment”. In: Cancer Communications 42.11 (2022), pp. 1083–1111.

[10] Valentin Gensbittel et al. “Mechanical adaptability of tumor cells in metastasis”. In: Developmental cell 56.2 (2021), pp. 164–179.

[11] FuiBoon Kai, Allison P Drain, and Valerie M Weaver. “The extracellular matrix modulates the metastatic journey”. In: Developmental cell 49.3 (2019), pp. 332–346.

[12] Charlotte Alibert, Bruno Goud, and Jean-Baptiste Manneville. “Are cancer cells really softer than normal cells?” In: Biology of the Cell 109.5 (2017), pp. 167–189.

[13] Douglas Hanahan. “Hallmarks of cancer: new dimensions”. In: Cancer discovery 12.1 (2022), pp. 31–46.

[14] Kin-Hoe Chow, Rachel E Factor, and Katharine S Ullman. “The nuclear envelope environment and its cancer connections”. In: Nature Reviews Cancer 12.3 (2012), pp. 196–209.

[15] H Martin et al. “Automated image analysis of gliomas an objective and reproducible method for tumor grading”. In: Acta neuropathologica 63 (1984), pp. 160–169.

[16] Katarzyna Pogoda et al. “Compression stiffening of brain and its effect on mechanosensing by glioma cells”. In: New journal of physics 16.7 (2014), p. 075002.

[17] Thomas James Grundy et al. “Differential response of patient-derived primary glioblastoma cells to environmental stiffness”. In: Scientific reports 6.1 (2016), p. 23353.

[18] J Matthew Barnes et al. “A tension-mediated glycocalyx–integrin feedback loop promotes mesenchymal-like glioblastoma”. In: Nature cell biology 20.10 (2018), pp. 1203–1214.

[19] Ariane E Erickson et al. “Fabrication and characterization of chitosan–hyaluronic acid scaffolds with varying stiffness for glioblastoma cell culture”. In: Advanced healthcare materials 7.15 (2018), p. 1800295.

[20] Celine Denais and Jan Lammerding. “Nuclear mechanics in cancer”. In: Cancer biology and the nuclear envelope: Recent advances may elucidate past paradoxes (2014), pp. 435–470.

[21] Claudia Tanja Mierke. “The fundamental role of mechanical properties in the progression of cancer disease and inflammation”. In: Reports on Progress in Physics 77.7 (2014), p. 076602.

[22] Claudia Tanja Mierke. “The matrix environmental and cell mechanical properties regulate cell migration and contribute to the invasive phenotype of cancer cells”. In: Reports on Progress in Physics 82.6 (2019), p. 064602.

[23] Yosef Gruenbaum and Roland Foisner. “Lamins: nuclear intermediate filament proteins with fundamental functions in nuclear mechanics and genome regulation”. In: Annual review of biochemistry 84.1 (2015), pp. 131–164.

[24] Youngjo Kim. “The impact of altered lamin B1 levels on nuclear lamina structure and function in aging and human diseases”. In: Current Opinion in Cell Biology 85 (2023), p. 102257. issn: 0955-0674. doi: 10.1016/j.ceb.2023.102257. url: https://www.sciencedirect.com/science/article/pii/S0955067423001060.

[25] Robert D Goldman et al. “Nuclear lamins: building blocks of nuclear architecture”. In: Genes & development 16.5 (2002), pp. 533–547.

[26] Thomas Dechat et al. “Nuclear lamins”. In: Cold Spring Harbor perspectives in biology 2.11 (2010), a000547.

[27] Amnon Buxboim et al. “Scaffold, mechanics and functions of nuclear lamins”. In: FEBS letters 597.22 (2023), pp. 2791–2805.

[28] Howard J Worman. “Nuclear lamins and laminopathies”. In: The Journal of pathology 226.2 (2012), pp. 316–325.

[29] Niina Dubik and Sabine Mai. “Lamin A/C: function in normal and tumor cells”. In: Cancers 12.12 (2020), p. 3688.

[30] Joe Swift et al. “Nuclear lamin-A scales with tissue stiffness and enhances matrix-directed differentiation”. In: Science 341.6149 (2013), p. 1240104.

[31] Celine M Denais et al. “Nuclear envelope rupture and repair during cancer cell migration”. In: Science 352.6283 (2016), pp. 353–358.

[32] Yuan Xie et al. “The human glioblastoma cell culture resource: validated cell models representing all molecular subtypes”. In: EBioMedicine 2.10 (2015), pp. 1351–1363.

[33] Laure Coquand et al. “A cell fate decision map reveals abundant direct neurogenesis bypassing intermediate progenitors in the human developing neocortex”. In: Nature Cell Biology 26.5 (May 2024), pp. 698–709. issn: 1476-4679. doi: 10.1038/s41556-024-01393-z. url: 10.1038/s41556-024-01393-z.

[34] David Guet et al. “Mechanical role of actin dynamics in the rheology of the Golgi complex and in Golgi-associated trafficking events”. In: Current Biology 24.15 (2014), pp. 1700–1711.

[35] Charlotte Alibert et al. “Multiscale rheology of glioma cells”. In: Biomaterials 275 (2021), p. 120903.

[36] Jean-Yves Tinevez et al. “TrackMate: An open and extensible platform for single-particle tracking”. In: Methods 115 (2017), pp. 80–90.

[37] Sandrine Etienne-Manneville, Florent Peglion, and Franck Coumailleau. “Live Imaging of Microtubule Dynamics in Glioblastoma Cells Invading the Zebrafish Brain”. In: JoVE 185 (July 2022). Publisher: MyJoVE Corp, e64093. issn: 1940-087X. doi: 10.3791/64093. url: https://app.jove.com/64093.

[38] Emma J van Bodegraven et al. “Intermediate filaments promote glioblastoma cell invasion by controlling cell deformability and mechanosensitive gene expression”. In: (2023). doi: 10.21203/rs.3.rs-2828066/v1.

[39] Takeshi Shimi et al. “The A-and B-type nuclear lamin networks: microdomains involved in chromatin organization and transcription”. In: Genes & development 22.24 (2008), pp. 3409–3421.

[40] Bruce Nmezi et al. “Concentric organization of A-and B-type lamins predicts their distinct roles in the spatial organization and stability of the nuclear lamina”. In: Proceedings of the National Academy of Sciences 116.10 (2019), pp. 4307–4315.

[41] Takeshi Shimi et al. “Structural organization of nuclear lamins A, C, B1, and B2 revealed by superresolution microscopy”. In: Molecular biology of the cell 26.22 (2015), pp. 4075–4086.

[42] Amir Vahabikashi et al. “Nuclear lamin isoforms differentially contribute to LINC complex-dependent nucleocytoskeletal coupling and whole-cell mechanics”. In: Proceedings of the National Academy of Sciences 119.17 (2022), e2121816119.

[43] Jan Lammerding et al. “Lamin A/C deficiency causes defective nuclear mechanics and mechanotransduction”. In: The Journal of clinical investigation 113.3 (2004), pp. 370–378.

[44] Patricia M Davidson and Jan Lammerding. “Broken nuclei–lamins, nuclear mechanics, and disease”. In: Trends in cell biology 24.4 (2014), pp. 247–256.

[45] Jan Lammerding et al. “Lamins A and C but not lamin B1 regulate nuclear mechanics”. In: Journal of Biological Chemistry 281.35 (2006), pp. 25768–25780.

[46] Nana Naetar, Simona Ferraioli, and Roland Foisner. “Lamins in the nuclear interior- life outside the lamina”. In: Journal of cell science 130.13 (2017), pp. 2087–2096.

[47] Natalie Y Chen et al. “An absence of lamin B1 in migrating neurons causes nuclear membrane ruptures and cell death”. In: Proceedings of the National Academy of Sciences 116.51 (2019), pp. 25870–25879.

[48] Olga Moiseeva et al. “Retinoblastoma-independent regulation of cell proliferation and senescence by the p53–p21 axis in lamin A/C-depleted cells”. In: Aging Cell 10.5 (2011), pp. 789–797. doi: 10.1111/j.1474-9726.2011.00719.x.

[49] T Sullivan et al. “Loss of A-type lamin expression compromises nuclear envelope integrity leading to muscular dystrophy”. en. In: J. Cell Biol. 147.5 (Nov. 1999), pp. 913–920.

[50] Vicente Andrés and José M González. “Role of A-type lamins in signaling, transcription, and chromatin organization”. In: Journal of Cell Biology 187.7 (2009), pp. 945–957.

[51] Josef Gotzmann and Roland Foisner. “Lamins and lamin-binding proteins in functional chromatin organizationr”. In: Critical Reviews™ in Eukaryotic Gene Expression 9.3-4 (1999).

[52] Cécile Leduc and Sandrine Etienne-Manneville. “Intermediate filaments in cell migration and invasion: the unusual suspects”. In: Current opinion in cell biology 32 (2015), pp. 102–112.

[53] Guilherme Pedreira de Freitas Nader et al. “Compromised nuclear envelope integrity drives TREX1-dependent DNA damage and tumor cell invasion”. In: Cell 184.20 (2021), 5230–5246.e22. issn: 0092-8674. doi: 10.1016/j.cell.2021.08.035. url: https://www.sciencedirect.com/science/article/pii/S0092867421010461.

[54] Tony Fischer, Alexander Hayn, and Claudia Tanja Mierke. “Effect of Nuclear Stiffness on Cell Mechanics and Migration of Human Breast Cancer Cells”. In: Frontiers in Cell and Developmental Biology 8 (2020). issn: 2296-634X. doi: 10.3389/fcell.2020.00393. url: https://www.frontiersin.org/journals/cell-and-developmental-biology/articles/10.3389/fcell.2020.00393.

[55] Takamasa Harada et al. “Nuclear lamin stiffness is a barrier to 3D migration, but softness can limit survival”. In: Journal of Cell Biology 204.5 (2014), pp. 669–682.

[56] Carlos F Guimarães et al. “The stiffness of living tissues and its implications for tissue engineeringung”. In: Nature Reviews Materials 5.5 (2020), pp. 351–370.

[57] Giorgio Seano and Rakesh K Jain. “Vessel co-option in glioblastoma: emerging insights and opportunities”. In: Angiogenesis 23.1 (2020), pp. 9–16.

[58] Rajesh Kumar Gupta et al. “Tumor-specific migration routes of xenotransplanted human glioblastoma cells in mouse brain”. In: Scientific Reports 14.1 (2024), p. 864.

[59] Sergej Skvortsov et al. “Proteomics profiling of microdissected low-and high-grade prostate tumors identifies Lamin A as a discriminatory biomarker”. In: Journal of proteome research 10.1 (2011), pp. 259–268.

[60] EJ Th Belt et al. “Loss of lamin A/C expression in stage II and III colon cancer is associated with disease recurrence”. In: European journal of cancer 47.12 (2011), pp. 1837–1845.

[61] SF Moss et al. “Decreased and aberrant nuclear lamin expression in gastrointestinal tract neoplasms”. In: Gut 45.5 (1999), pp. 723–729.

[62] Zhengrong Wu et al. “Reduced expression of lamin A/C correlates with poor histological differentiation and prognosis in primary gastric carcinoma”. In: Journal of Experimental & Clinical Cancer Research 28 (2009), pp. 1–12.

[63] Callinice D Capo-chichi et al. “Loss of A-type lamin expression compromises nuclear envelope integrity in breast cancer”. In: Chinese journal of cancer 30.6 (2011), p. 415.

[64] Giovanna Maresca et al. “LMNA knock-down affects differentiation and progression of human neuroblastoma cells”. In: (2012).

[65] Kunnathur Murugesan Sakthivel and Poonam Sehgal. “A novel role of lamins from genetic disease to cancer biomarkers”. In: Oncology reviews 10.2 (2016).

[66] Naomi D Willis et al. “Lamin A/C is a risk biomarker in colorectal cancer”. In: PloS one 3.8 (2008), e2988.

[67] Hea-Jin Jung et al. “Nuclear lamins in the brain—new insights into function and regulation”. In: Molecular neurobiology 47 (2013), pp. 290–301.

[68] Hea-Jin Jung et al. “Regulation of prelamin A but not lamin C by miR-9, a brain-specific microRNA”. In: Proceedings of the National Academy of Sciences 109.7 (2012), E423–E431.

[69] Brian C Capell and Francis S Collins. “Human laminopathies: nuclei gone genetically awry”. In: Nature reviews genetics 7.12 (2006), pp. 940–952.

[70] David A Parry et al. “Heterozygous lamin B1 and lamin B2 variants cause primary microcephaly and define a novel laminopathy”. In: Genetics in Medicine 23.2 (2021), pp. 408–414.

