## Supplementary document for "Lamins regulate nuclear mechanics and shape to control glioblastoma cell proliferation, migration and invasion"

### Supplementary Information

#### Supplementary Methods

#### Supplementary Figures

#### Supplementary Movies

#### Supplementary Tables

### Supplementary Methods

Table 1: **Antibodies used for immunofluorescence and Western Blot experiments.**

| Antibody | Product code | Company | Application | Species |
|---|---|---|---|---|
| LAMIN A/C | SAB4200236 2 | Abcam | IF, WB | Mouse |
| LAMIN B1 | ab16048 | Abcam | IF, WB | Rabbit |
| $\beta$-actin | 66009 | Proteinteck | WB | Mouse |

**Table of antibodies**

**Track Mate parameters** Parameter used in TrackMate plugin to analyze the Iprasense recorded images:

Table 2: **Trackmate parameters.**

| Parameters | Values |
|---|---|
| Pixel | [20, 25] |
| Frame linking distance | [30, 45] |
| Track segment gap distance | [30, 45] |
| Max frame gap | 3 |

**Nuclear shape index definitions:**

$$Circularity = \frac{4\pi * area}{perimeter^2}$$

$$ShapeIndex = \frac{perimeter}{2\sqrt{area\pi}}$$

$$Roundness = \frac{4area}{\pi * majoraxis^2}$$

### Supplementary Figures

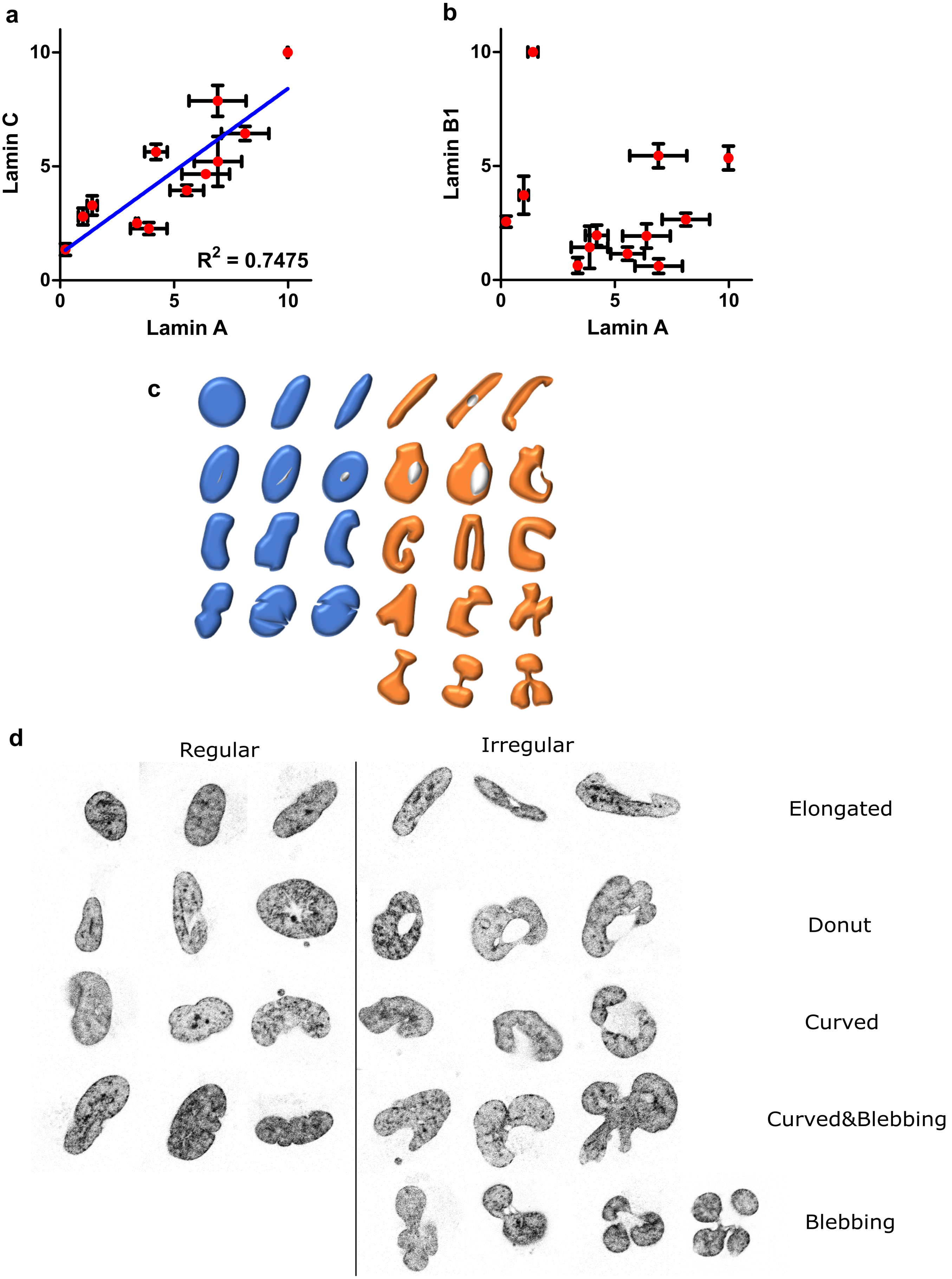

a
Lamin C
Lamin A
R2 = 0.7475
b
Lamin B1
Lamin A
c
d
Regular
Irregular
Elongated
Donut
Curved
Curved&Blebbing
Blebbing

**Supplementary Figure** 1: **Lamin expression levels and nuclear morphology. a** Correlation between the expression levels of lamin C and lamin A. A positive correlation is observed (linear fit). **b** Correlation between the expression levels of lamin B1 and lamin A. No correlation is observed. **c** Classification of nuclear shapes as regular (blue) and irregular (orange) nuclei. **d** Images of nuclei stained with Hoechst in GBM cell lines and in RG cells. Four subgroups of irregular nuclei are distinguished: elongated, donut, curved, and blebbing.

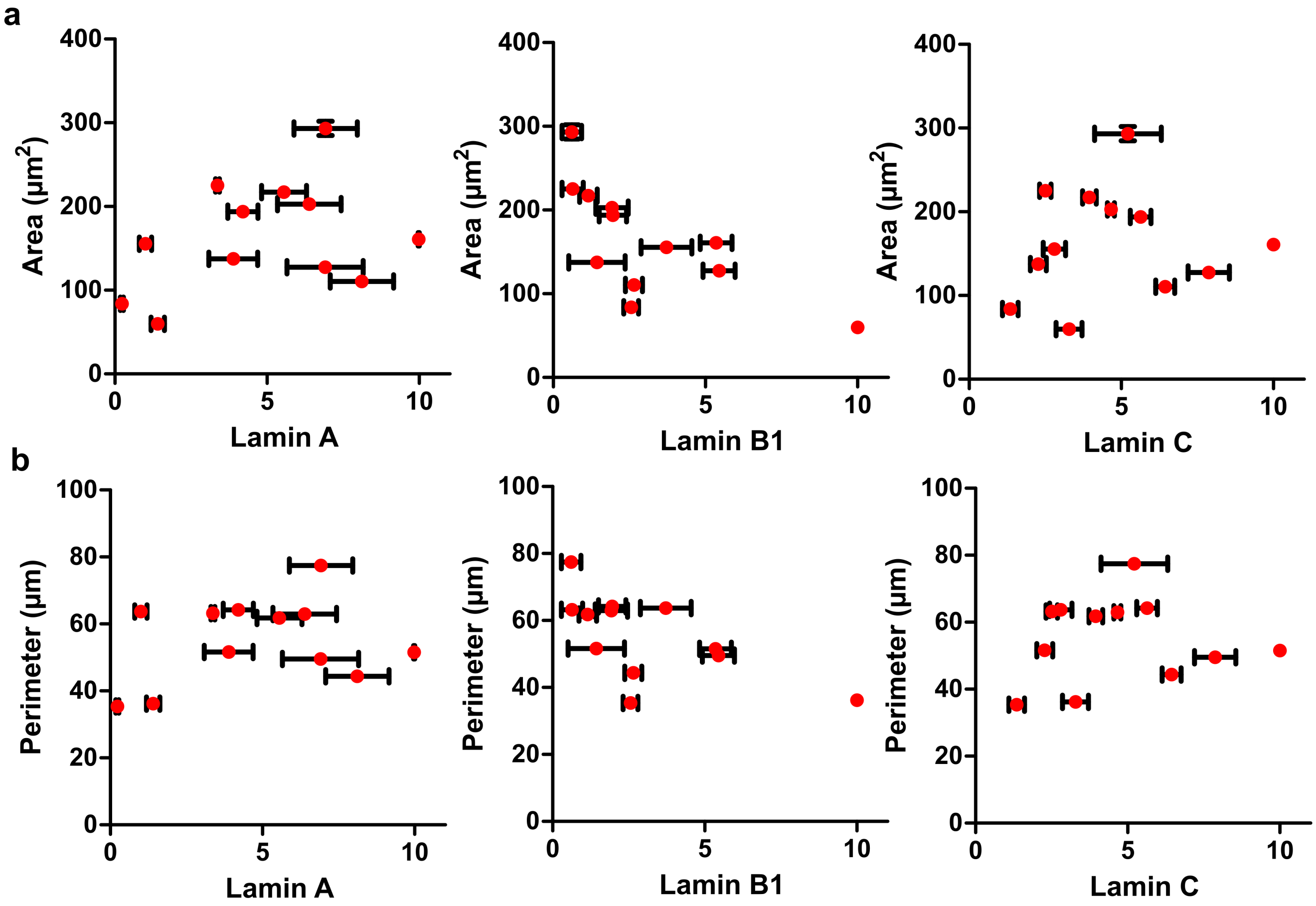


**Supplementary Figure** 2: **Lamin expression levels and nuclear size. a** Correlation between the nuclear projected area and the expression levels of lamin A, lamin B1, and lamin C. **b** Correlation between the nuclear projected perimeter and the expression levels of lamin A, lamin B1, and lamin C.

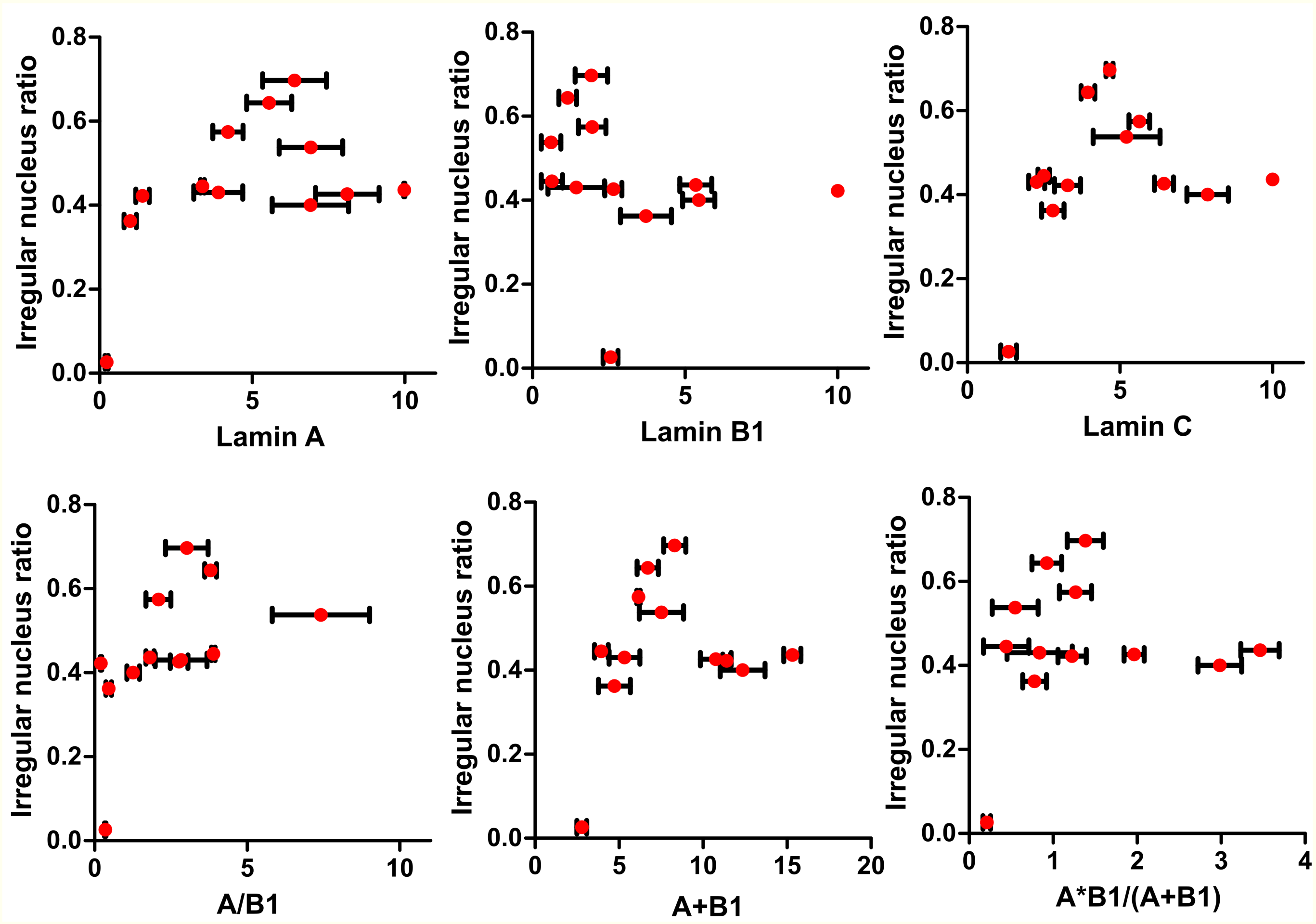


**Supplementary Figure** 3: **Lamin expression levels and proportion of irregular nuclei. a** Correlation between the proportion of irregular nuclei and the expression levels of lamin A, lamin B1, and lamin C. **b** Correlation between the proportion of irregular nuclei and the quantities A/B1, A+B1 and (A*B1)/(A+B1) where A is the expression level of lamin A and B1 is the expression level of lamin B1.

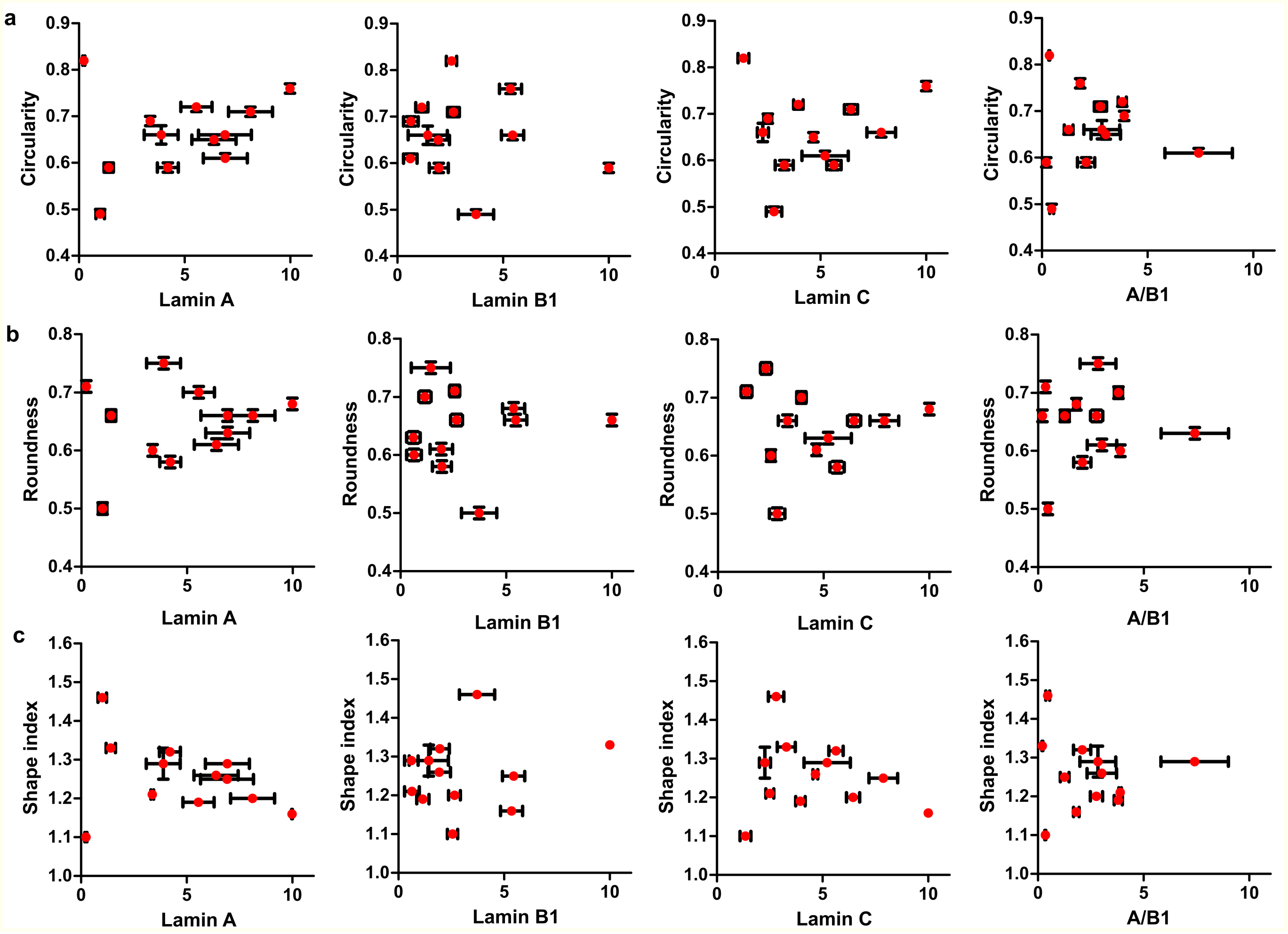


**Supplementary Figure** 4: **Lamin expression levels and nuclear morphological parameters.** Correlation between nuclear circularity **a**, roundness **b** and shape index **c** and the expression levels of lamin A, lamin B1, and lamin C, and the lamin ratio A/B1.

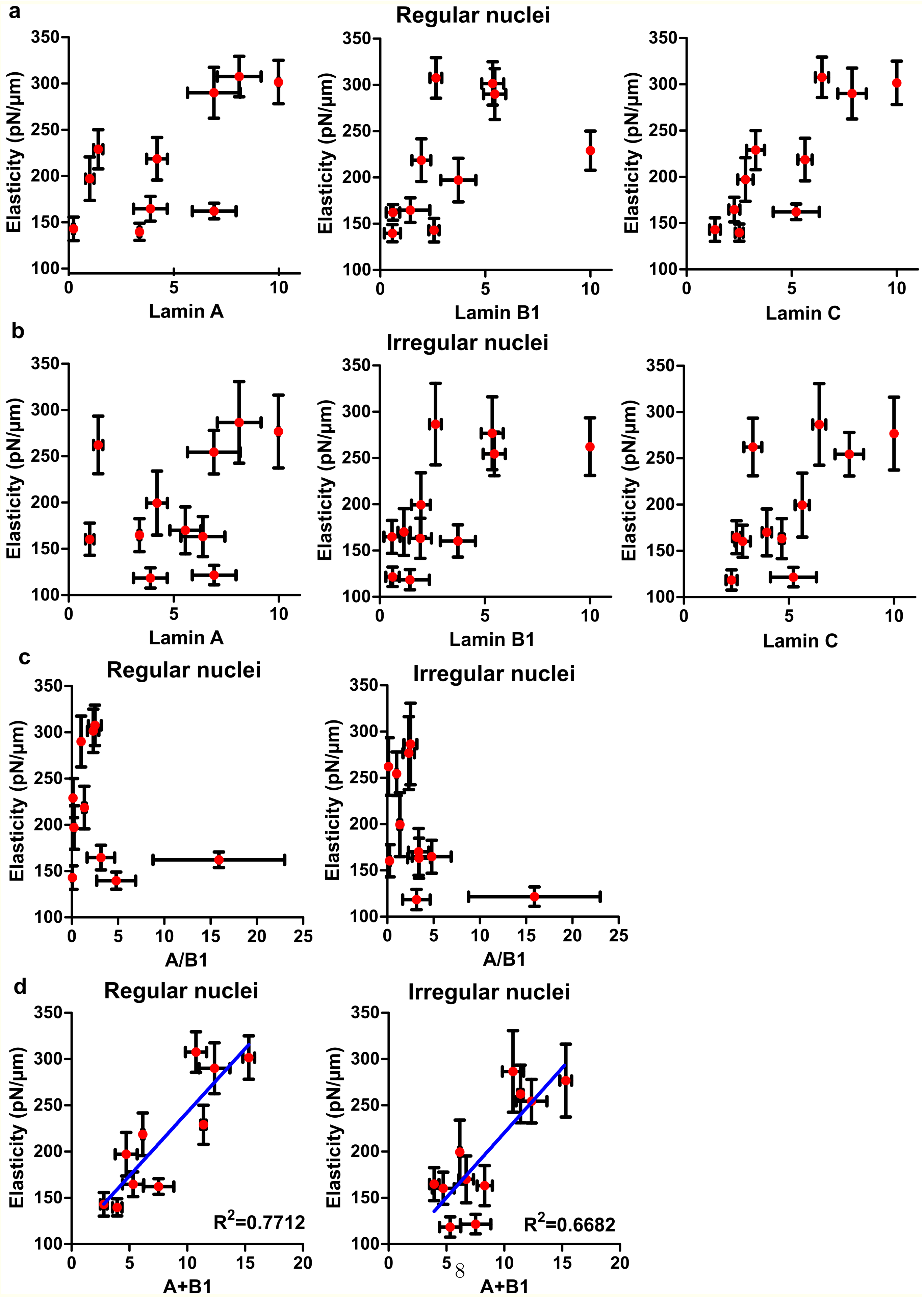
a
Regular nuclei
Elasticity (pN/µm)
Lamin A
Lamin B1
Lamin C
b
Irregular nuclei
Elasticity (pN/µm)
Lamin A
Lamin B1
Lamin C
c
Regular nuclei
Irregular nuclei
Elasticity (pN/µm)
A/B1
d
Regular nuclei
Irregular nuclei
Elasticity (pN/µm)
$R^2$=0.7712
$R^2$=0.6682
A+B1

**Supplementary Figure** 5: **Lamin expression levels and nuclear elasticity. a-b** Correlation between the elasticity of regular (a) and irregular (b) nuclei and the expression levels of lamin A, lamin B1, and lamin C. **c** Correlation between the elasticity of regular (left) and irregular (right) nuclei and the lamin ratio A/B1. **d** Correlation between the elasticity of regular (left) and irregular (right) nuclei and the sum of lamin A and lamin B1 expression levels denoted as A+B1. Spearman $r^2$ values are indicated.

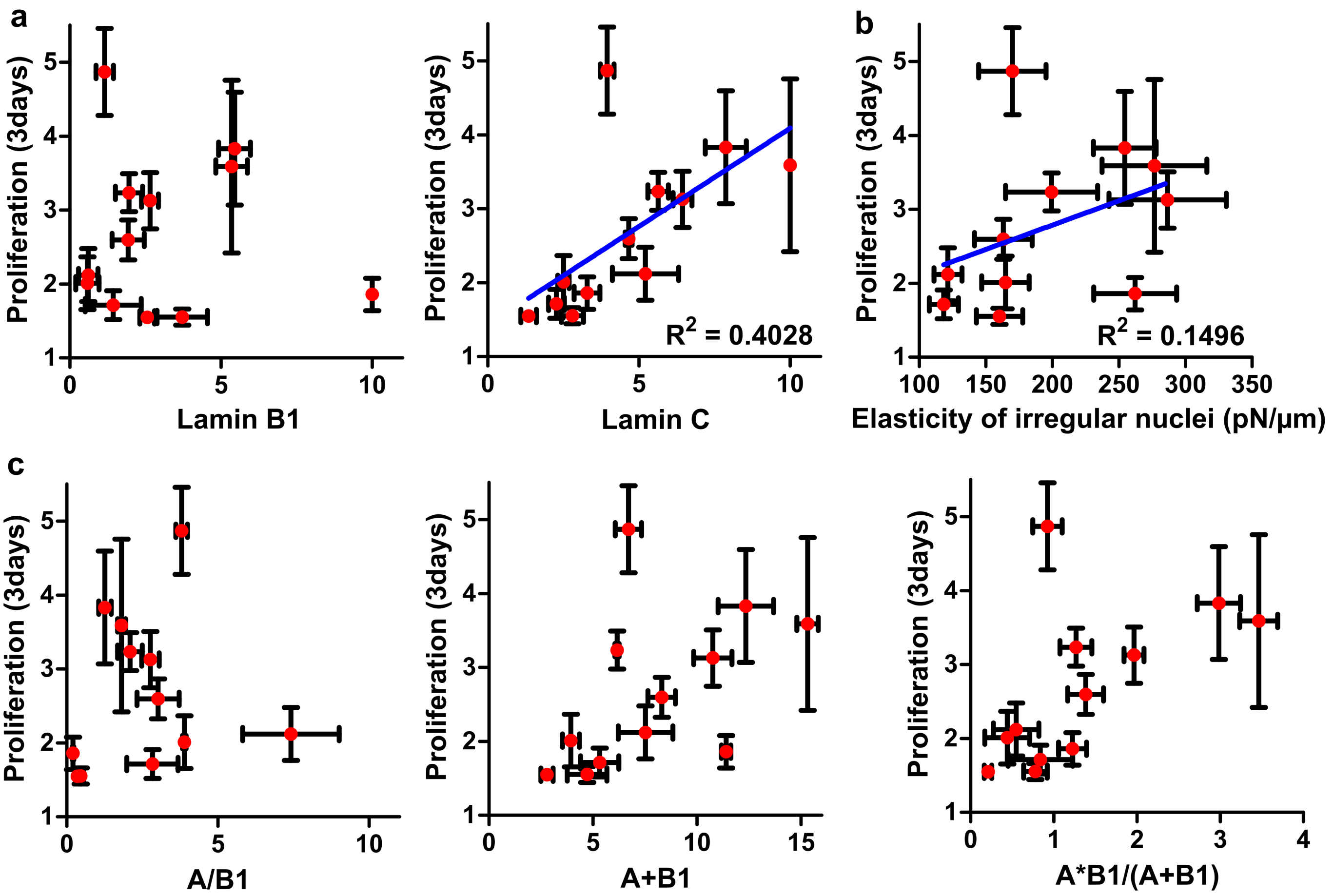


**Supplementary Figure** 6: **Lamin expression levels and cell proliferation. a** Correlation between proliferation rates and the expression levels of lamin B1 and lamin C. **b** Correlation between proliferation rates and the elasticity of irregular nuclei. **c** Correlation between proliferation rates and the quantities A/B1, A+B1 and (A*B1)/(A+B1) where A is the expression level of lamin A and B1 is the expression level of lamin B1.

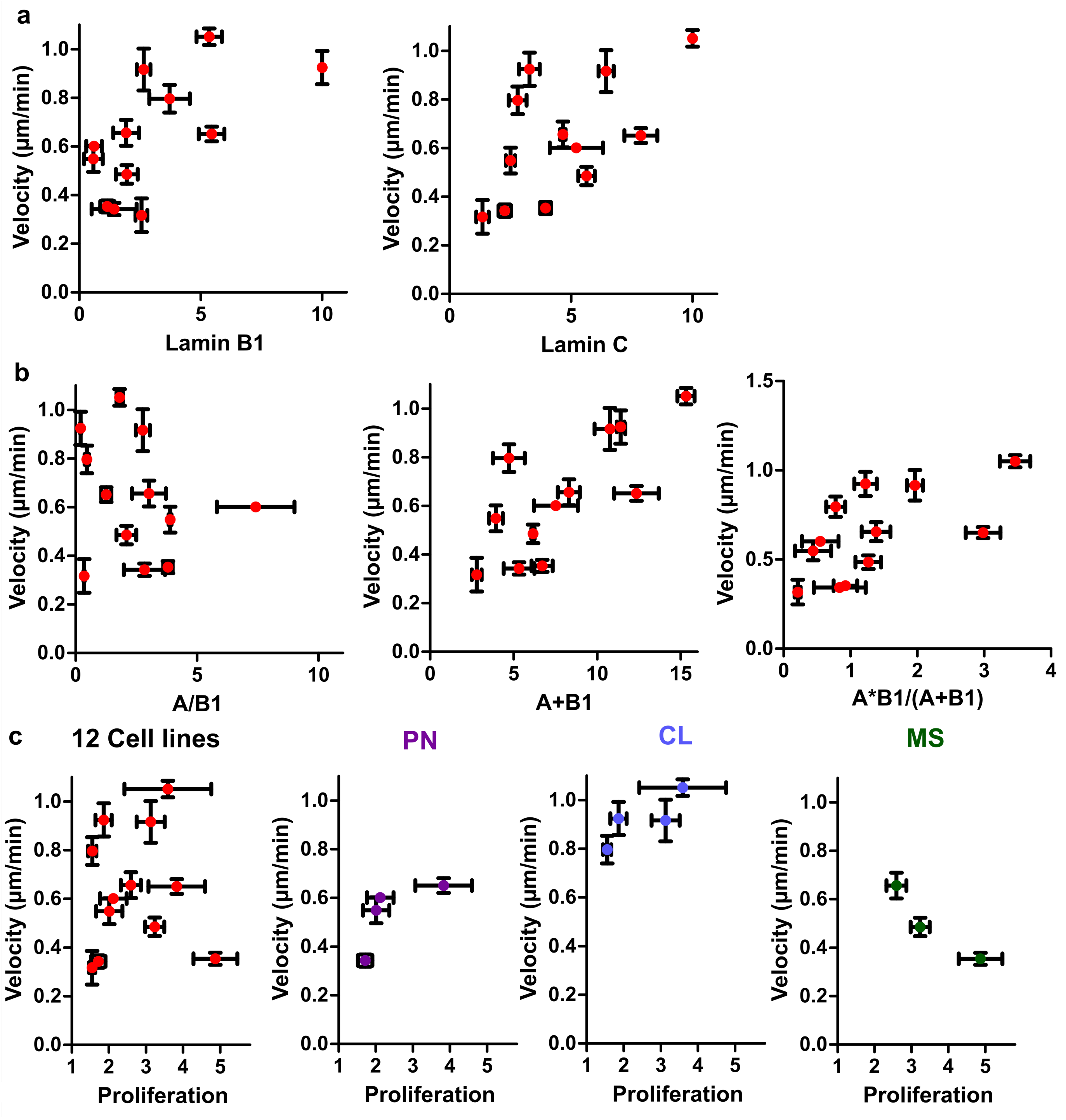


**Supplementary Figure** 7: **Lamin expression levels and 2D cell migration. a** Correlation between the 2D migration velocity and the expression levels of lamin B1 and lamin C. **b** Correlation between the 2D migration velocity and the quantities A/B1, A+B1 and (A*B1)/(A+B1) where A is the expression level of lamin A and B1 is the expression level of lamin B1. **c** Correlation between the 2D migration velocity and the proliferation rate for the twelve cell lines (left panel), and for the three different phenotypes proneural (PN), classical (CL), mesenchymal (MS) plotted separately.

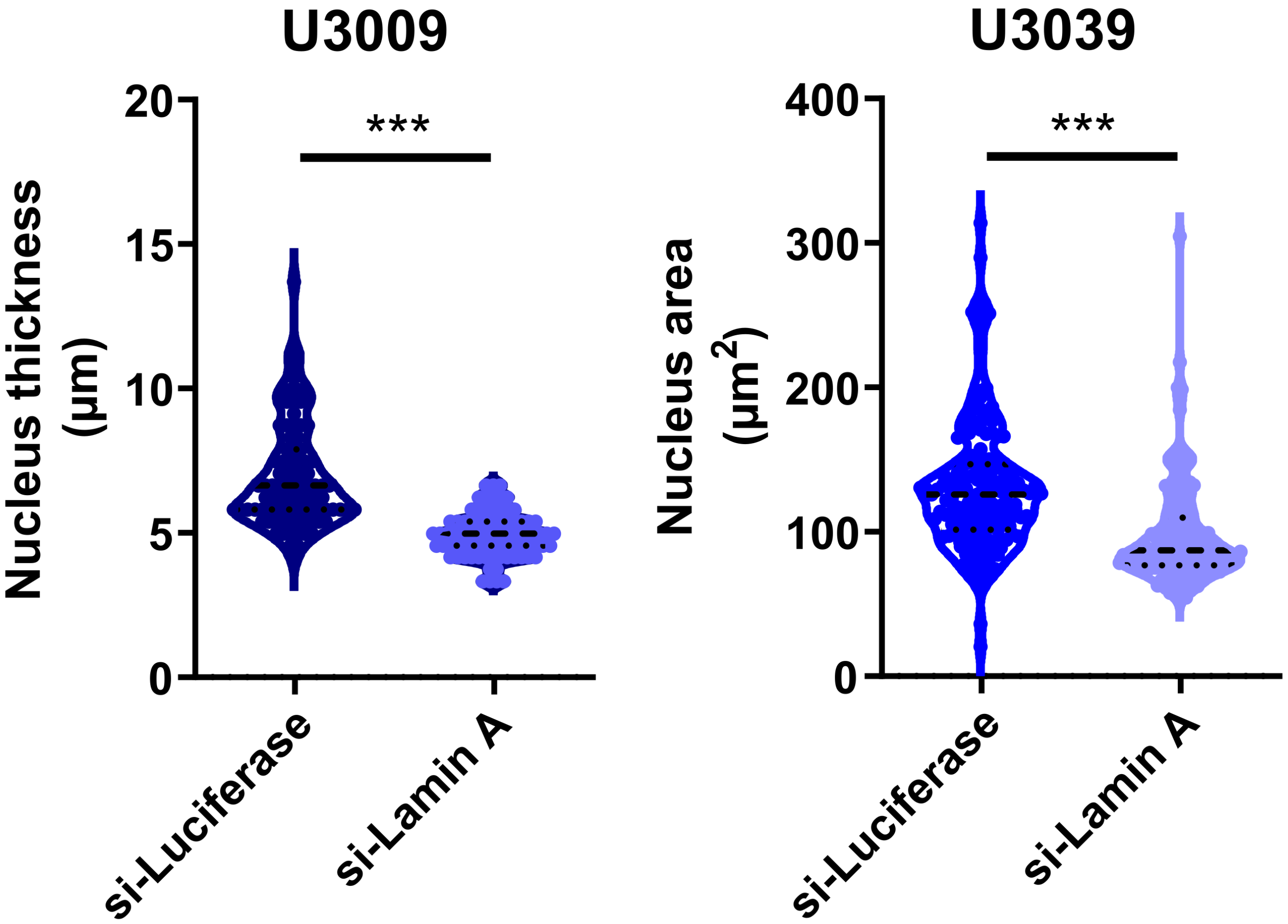


**Supplementary Figure** 8: **Effect of lamin A depletion on nuclear morphology.** (Left) The thickness of the nucleus in U3009 cells treated with si-Lamin A decreases compared to control cells treated with si-Luciferase. (Right) The 2D projected area of the nucleus in U3039 cells treated with si-Lamin A decreases compared to control cells treated with si-Luciferase.

#### Supplementary movies

**Supplementary Movie 1** Typical indentation experiment of a regular nucleus. The 2µm-diameter bead trapped by optical tweezers is shown in white. The nucleus stained with Hoechst is shown in blue. The stage is moved downwards to indent the nucleus. Corresponding still images are shown in Fig. 2a (upper panel). Scale bar, 2 µm. Real-time duration, 64 seconds

**Supplementary Movie 2** Typical indentation experiment of an irregular nucleus. The 2µm-diameter bead trapped by optical tweezers is shown in white. The nucleus stained with Hoechst is shown in blue. The stage is moved downwards to indent the nucleus. Corresponding still images are shown in Fig. 2b (lower panel). Scale bar, 2 µm. Real-time duration, 64 seconds.

**Supplementary Movie 3** Lens-free microscopy imaging to quantify cell migration and proliferation during 72 hours. Scale bar, 500 µm.

### Supplementary Tables

| Correlation between lamin C and lamin A | |
|---|---|
| Linear | Elasticity (pN/$\mu$m) |
| Slope | 0.7258 ± 0.1334 |
| Y-intercept when X=0.0 | 1.159 ± 0.7504 |
| X-intercept when Y=0.0 | -1.597 |
| 1/slope | 1.378 |
| 95% Confidence Intervals | |
| Slope | 0.4286 to 1.023 |
| Y-intercept when X=0.0 | -0.5125 to 2.831 |
| X-intercept when Y=0.0 | -6.308 to 0.5246 |
| Goodness of Fit | |
| R square | 0.7475 |
| Sy.x | 1.333 |
| Is slope significantly non-zero? | |
| F | 29.61 |
| DFn. DFd | 1.000, 10.000 |
| P value | 0.0003 |
| Deviation from zero? | Significant |
| Data | |
| Number of X values | 12 |
| Maximum number of Y replicates | 1 |
| Total number of values | 12 |
| Number of missing values | 0 |

Table 3: Correlation between lamin C and lamin A

| Correlation between nuclear area and lamin ratio A/B1 | |
|---|---|
| One phase decay: Best-fit values | Nuclei area($\mu m^2$) |
| $Y_{min}$ | 97.59 |
| $Y_{max}$ | 311.4 |
| k | 0.1588 |
| Std. Error | |
| $Y_{min}$ | 22.52 |
| $Y_{max}$ | 64.5 |
| k | 0.09714 |
| 95% Confidence Intervals | |
| $Y_{min}$ | 46.66 to 148.5 |
| $Y_{max}$ | 165.5 to 457.2 |
| k | 0.0 to 0.3786 |
| Goodness of Fit | |
| Degrees of Freedom | 9 |
| R square | 0.7041 |
| Absolute Sum of Squares | 14207 |
| Sy.x | 39.73 |
| Constraints | |
| k | k > 0.0 |
| Number of points | |
| Analyzed | 12 |

Table 4: Correlation of nuclear area and lamin ratio A/B1

| Correlation between nuclear perimeter and lamin ratio A/B1 | |
|---|---|
| One phase decay: Best-fit values | Nuclei perimeter($\mu$m) |
| $Y_{min}$<br>$Y_{max}$<br>k | 45.14<br>82.09<br>0.1293 |
| Std. Error | |
| $Y_{min}$<br>$Y_{max}$<br>k | 5.496<br>20.47<br>0.1402 |
| 95% Confidence Intervals | |
| $Y_{min}$<br>$Y_{max}$<br>k | 32.71 to 57.58<br>35.79 to 128.4<br>0.0 to 0.4465 |
| Goodness of Fit | |
| Degrees of Freedom<br>R square<br>Absolute Sum of Squares<br>Sy.x | 9<br>0.5025<br>867.8<br>9.82 |
| Constraints | |
| k | k > 0.0 |
| Number of points | |
| Analyzed | 12 |

Table 5: Correlation of nuclear perimeter and lamin ratio A/B1

| Correlation between regular nuclei elasticity and the sum (A+B1) | |
|---|---|
| Linear | Elasticity (pN/$\mu$m) |
| Slope | 13.74 ± 2.646 |
| Y-intercept when X=0.0 | 105.0 ± 23.70 |
| X-intercept when Y=0.0 | -7.645 |
| 1/slope | 0.07278 |
| 95% Confidence Intervals | |
| Slope | 7.639 to 19.84 |
| Y-intercept when X=0.0 | 50.41 to 159.7 |
| X-intercept when Y=0.0 | -20.39 to -2.605 |
| Goodness of Fit | |
| R square | 0.7712 |
| Sy.x | 33.14 |
| Is slope significantly non-zero? | |
| F | 26.97 |
| DFn. DFd | 1.000. 8.000 |
| P value | 0.0008 |
| Deviation from zero? | Significant |
| Data | |
| Number of X values | 10 |
| Maximum number of Y replicates | 1 |
| Total number of values | 10 |
| Number of missing values | 2 |

Table 6: Correlation between regular nuclei elasticity and the sum of the expression levels of lamin A and lamin B1 (A+B1)

| Correlation between irregular nuclei elasticity and the sum (A+B1) | |
|---|---|
| Linear | Elasticity (pN/$\mu$m) |
| Slope | 14.00 ± 3.288 |
| Y-intercept when X=0.0 | 80.26 ± 29.87 |
| X-intercept when Y=0.0 | -5.734 |
| 1/slope | 0.07145 |
| 95% Confidence Intervals | |
| Slope | 6.560 to 21.43 |
| Y-intercept when X=0.0 | 12.68 to 147.8 |
| X-intercept when Y=0.0 | -21.99 to -0.6064 |
| Goodness of Fit | |
| R square | 0.6682 |
| Sy.x | 37.44 |
| Is slope significantly non-zero? | |
| F | 18.13 |
| DFn. DFd | 1.000, 9.000 |
| P value | 0.0021 |
| Deviation from zero? | Significant |
| Data | |
| Number of X values | 11 |
| Maximum number of Y replicates | 1 |
| Total number of values | 11 |
| Number of missing values | 1 |

Table 7: Correlation between irregular nuclei elasticity and the sum of the expression levels of lamin A and lamin B1

| Correlation between regular nuclei elasticity and the effective lamin level L=(A*B1)/(A+B1) | |
|---|---|
| Plateau followed by one phase decay | Elasticity (pN/$\mu$m) |
| Best-fit values | |
| $L_0$ | 0.4834 |
| $Y_{min}$ | 141.4 |
| $Y_{max}$ | 316.5 |
| $a$ | 0.9506 |
| Std. Error | |
| $L_0$ | 0.1515 |
| $Y_{min}$ | 14.4 |
| $Y_{max}$ | 26.61 |
| $a$ | 0.4265 |
| 95% Confidence Intervals | |
| $L_0$ | 0.1128 to 0.8541 |
| $Y_{min}$ | 106.1 to 176.6 |
| $Y_{max}$ | 251.4 to 381.6 |
| $a$ | 0.0 to 1.994 |
| Goodness of Fit | |
| Degrees of Freedom | 6 |
| R square | 0.9352 |
| Absolute Sum of Squares | 2490 |
| Sy.x | 20.37 |
| Constraints | |
| $a$ | $a > 0.0$ |
| Number of points | |
| Analyzed | 10 |

Table 8: Correlation between regular nuclei elasticity and the effective lamin level L=(A*B1)/(A+B1)

| Correlation between irregular nuclei elasticity and the effective lamin level L=(A*B1)/(A+B1) | |
|---|---|
| Plateau followed by one phase decay | Elasticity (pN/$\mu$m) |
| Best-fit values | |
| $L_0$ | 0.8389 |
| $Y_{min}$ | 144.1 |
| $Y_{max}$ | 274.1 |
| $a$ | 1.689 |
| Std. Error | |
| $L_0$ | 0.2349 |
| $Y_{min}$ | 21.17 |
| $Y_{max}$ | 31.05 |
| $a$ | 1.434 |
| 95% Confidence Intervals | |
| $L_0$ | 0.2834 to 1.395 |
| $Y_{min}$ | 94.05 to 194.2 |
| $Y_{max}$ | 200.7 to 347.5 |
| $a$ | 0.0 to 5.081 |
| Goodness of Fit | |
| Degrees of Freedom | 7 |
| R square | 0.7383 |
| Absolute Sum of Squares | 9949 |
| Sy.x | 37.7 |
| Constraints | |
| $a$ | $a > 0.0$ |
| Number of points | |
| Analyzed | 11 |

Table 9: Correlation between irregular nuclei elasticity and the effective lamin level L=(A*B1)/(A+B1)

| Correlation between proliferation and lamin A expression | |
|---|---|
| Linear | Elasticity (pN/$\mu$m) |
| Slope | 0.2271 $\pm$ 0.08500 |
| Y-intercept when X=0.0 | 1.574 $\pm$ 0.4782 |
| X-intercept when Y=0.0 | -6.93 |
| 1/slope | 4.403 |
| 95% Confidence Intervals | |
| Slope | 0.03775 to 0.4165 |
| Y-intercept when X=0.0 | 0.5087 to 2.639 |
| X-intercept when Y=0.0 | -66.22 to -1.289 |
| Goodness of Fit | |
| R square | 0.4166 |
| Sy.x | 0.8493 |
| Is slope significantly non-zero? | |
| F | 7.14 |
| DFn. DFd | 1.000, 10.000 |
| P value | 0.0234 |
| Deviation from zero? | Significant |
| Data | |
| Number of X values | 12 |
| Maximum number of Y replicates | 1 |
| Total number of values | 12 |
| Number of missing values | 0 |

Table 10: Correlation between proliferation and lamin A expression levels

| Correlation between proliferation and lamin C expression | |
|---|---|
| Linear | Elasticity (pN/$\mu$m) |
| Slope | 0.2660 ± 0.1024 |
| Y-intercept when X=0.0 | 1,430 ± 0,5384 |
| X-intercept when Y=0.0 | -5.375 |
| 1/slope | 3.759 |
| 95% Confidence Intervals | |
| Slope | 0,03779 to 0,4943 |
| Y-intercept when X=0.0 | 0,2304 to 2,630 |
| X-intercept when Y=0.0 | -66,27 to -0,4895 |
| Goodness of Fit | |
| R square | 0.4028 |
| Sy.x | 0.4028 |
| Is slope significantly non-zero? | |
| F | 6.744 |
| DFn. DFd | 1.000, 10.000 |
| P value | 0.0266 |
| Deviation from zero? | Significant |
| Data | |
| Number of X values | 12 |
| Maximum number of Y replicates | 1 |
| Total number of values | 12 |
| Number of missing values | 0 |

Table 11: Correlation between proliferation and lamin C expression levels

| Correlation between proliferation and proportion of irregular nuclei | |
|---|---|
| Linear | Proliferation |
| Slope | 3.079 ± 1.731 |
| Y-intercept when X=0.0 | 1.284 ± 0.8281 |
| X-intercept when Y=0.0 | -0.4171 |
| 1/slope | 0.3247 |
| 95% Confidence Intervals | |
| Slope | -0.7775 to 6.936 |
| Y-intercept when X=0.0 | -0.5605 to 3.129 |
| X-intercept when Y=0.0 | -infinity to 0.08412 |
| Goodness of Fit | |
| R square | 0.2404 |
| Sy.x | 0.9691 |
| Is slope significantly non-zero? | |
| F | 3.164 |
| DFn. DFd | 1.000, 10.000 |
| P value | 0.1056 |
| Deviation from zero? | Not Significant |
| Data | |
| Number of X values | 12 |
| Maximum number of Y replicates | 1 |
| Total number of values | 12 |
| Number of missing values | 0 |

Table 12: Correlation between proliferation and proportion of irregular nuclei

| Correlation between proliferation and elasticity of regular nuclei | |
|---|---|
| Linear | Elasticity (pN/$\mu$m) |
| Slope | 0.01114 $\pm$ 0.002743 |
| Y-intercept when X=0.0 | 0.05894 $\pm$ 0.6149 |
| X-intercept when Y=0.0 | -5.291 |
| 1/slope | 89.76 |
| 95% Confidence Intervals | |
| Slope | 0.004815 to 0.01747 |
| Y-intercept when X=0.0 | -1.359 to 1.477 |
| X-intercept when Y=0.0 | -300.1 to 79.52 |
| Goodness of Fit | |
| R square | 0.6734 |
| Sy.x | 0.5377 |
| Is slope significantly non-zero? | |
| F | 16.49 |
| DFn. DFd | 1.000, 8.000 |
| P value | 0.0036 |
| Deviation from zero? | Significant |
| Data | |
| Number of X values | 10 |
| Maximum number of Y replicates | 1 |
| Total number of values | 10 |
| Number of missing values | 0 |

Table 13: Correlation between proliferation and elasticity of regular nuclei

| Correlation between proliferation and elasticity of irregular nuclei | |
|---|---|
| Linear | Elasticity (pN/$\mu$m) |
| Slope | 0.006576 ± 0.005227 |
| Y-intercept when X=0.0 | 1.471 ± 1.080 |
| X-intercept when Y=0.0 | -223.7 |
| 1/slope | 152.1 |
| 95% Confidence Intervals | |
| Slope | -0.005247 to 0.01840 |
| Y-intercept when X=0.0 | -0.9713 to 3.913 |
| X-intercept when Y=0.0 | -infinity to 54.75 |
| Goodness of Fit | |
| R square | 0.1496 |
| Sy.x | 1.019 |
| Is slope significantly non-zero? | |
| F | 1.583 |
| DFn. DFd | 1.000, 9.000 |
| P value | 0.24 |
| Deviation from zero? | Not Significant |
| Data | |
| Number of X values | 11 |
| Maximum number of Y replicates | 1 |
| Total number of values | 11 |

Table 14: Correlation between proliferation and elasticity of irregular nuclei

| Correlation between velocity and elasticity of regular nuclei | |
|---|---|
| Linear | Elasticity (pN/$\mu$m) |
| Slope | 0.002855 $\pm$ 0.0009171 |
| Y-intercept when X=0.0 | 0.04849 $\pm$ 0.2056 |
| X-intercept when Y=0.0 | -16.99 |
| 1/slope | 350.3 |
| 95% Confidence Intervals | |
| Slope | 0.0007398 to 0.004969 |
| Y-intercept when X=0.0 | -0.4255 to 0.5225 |
| X-intercept when Y=0.0 | -687.4 to 87.99 |
| Goodness of Fit | |
| R square | 0.5477 |
| Sy.x | 0.1797 |
| Is slope significantly non-zero? | |
| F | 9.689 |
| DFn. DFd | 1.000, 8.000 |
| P value | 0.0144 |
| Deviation from zero? | Significant |
| Data | |
| Number of X values | 10 |
| Maximum number of Y replicates | 1 |
| Total number of values | 10 |
| Number of missing values | 0 |

Table 15: Correlation between velocity and elasticity of regular nuclei

| Correlation between velocity and elasticity of irregular nuclei | |
|---|---|
| Linear | Elasticity (pN/$\mu$m) |
| Slope | 0.002806 ± 0.0008517 |
| Y-intercept when X=0.0 | 0.1105 ± 0.1759 |
| X-intercept when Y=0.0 | -39.37 |
| 1/slope | 356.4 |
| 95% Confidence Intervals | |
| Slope | 0.0008791 to 0.004732 |
| Y-intercept when X=0.0 | -0.2875 to 0.5084 |
| X-intercept when Y=0.0 | -564.6 to 62.23 |
| Goodness of Fit | |
| R square | 0.5466 |
| Sy.x | 0.1661 |
| Is slope significantly non-zero? | |
| F | 10.85 |
| DFn. DFd | 1.000, 9.000 |
| P value | 0.0093 |
| Deviation from zero? | Significant |
| Data | |
| Number of X values | 11 |
| Maximum number of Y replicates | 1 |
| Total number of values | 11 |
| Number of missing values | 0 |

Table 16: Correlation between velocity and elasticity of irregular nuclei
